# Dynamic landscapes of gene regulatory networks in early mammalian neurogenesis: Insights into brain evolution and disorder risk

**DOI:** 10.1101/2025.10.07.681016

**Authors:** Kalpana Hanthanan Arachchilage, Ryan D. Risgaard, Jie Sheng, Pubudu Kumarage, Jerome J. Choi, Sayali Anil Alatkar, Chirag Gupta, Xinyu Zhao, André M. M. Sousa, Daifeng Wang

## Abstract

Neurogenesis—the process of generating neurons—is governed by dynamic transcriptional programs that vary across time, brain regions, and cell types, forming regionally specialized neuronal circuits. To understand these dynamics, we constructed a comprehensive gene regulatory network (GRN) resource encompassing 22 neurogenic lineages from human, macaque, and mouse, enabling cross-species and cross-regional comparisons. Leveraging state-of-the-art trajectory analysis and GRN inference, we characterized temporal regulatory dynamics and introduced a “dynamic score” to identify key subnetworks with lineage-specific dynamics, including hundreds of regulons and co-regulatory modules. Our analysis uncovered both known and novel candidate regulators driving neuronal differentiation and regional identity, spanning the entire human brain, as well as evolutionary divergence in neurogenic GRNs distinguishing human brains. Mapping risk genes to the resource helped understand associated early gene regulatory dynamics with 35 neurodevelopmental disorders and traits including autism, schizophrenia, severe intellectual disability, and microcephaly. This resource is publicly available as an interactive online platform.

## INTRODUCTION

The evolution of the mammalian central nervous system (CNS) is characterized by a remarkable increase in the size, complexity, and functional specialization of discrete brain regions^1^. A growing body of evidence has revealed that major adaptations in brain structure and function are driven, in large part, by changes in gene regulation^2,3^. Differences in the timing, location, and magnitude of gene expression play a pivotal role in neural development. Comparative transcriptomic analyses have revealed cell-type-specific regulatory modifications, particularly within neuronal populations, that have contributed to the expansion of cortical surface area and emergence of novel connectivity patterns across the mammalian clade^4,5^. These gene regulatory differences orchestrate complex molecular programs during development and across cell types, driving the anatomical and cognitive diversification that distinguishes the mammalian, and particularly human, brain from those of other vertebrates.

Neurogenesis—a highly dynamic process fundamental to the formation and function of the CNS—is a critical developmental process during which evolutionary adaptation and diversification frequently shape neural architecture and complexity^6^. This intricate process encompasses the specification, differentiation, and maturation of neurons, each regulated by distinct transcriptional programs. Around 28-30 days post-fertilization (Carnegie stage 10), the developing human CNS undergoes significant patterning and regionalization. Morphogen gradients induce transcription factors (TFs) that define developing neural tube regions, culminating in the formation of the three primary vesicles: the prosencephalon (forebrain), mesencephalon (midbrain), and rhombencephalon (hindbrain). These give rise to distinct adult brain structures, including the telencephalon, diencephalon, midbrain, cerebellum, pons, and medulla oblongata^7^. Neuronal specification and regionalization establish the foundational organizational framework of the brain, enabling further subdivision and the development of specialized neural structures. The brain’s hierarchical anatomical structure is mirrored by a layered gene regulatory architecture, in which region-specific TFs initiate cascades of downstream regulatory interactions that progressively refine neuronal identity and organization^8,9^. These spatiotemporal regulatory dynamics ensure the precise patterning and differentiation of functionally diverse neuronal cell types.

Mapping gene regulatory relationships in the developing brain presents challenges not seen in conventional static gene regulatory networks (GRNs). Unlike systems with discrete transcriptomic states, brain development involves continuous, dynamic gene expression changes that require more nuanced modeling approaches. While numerous static, cell-type-specific regulatory network methods have been extensively applied in non-developmental settings^10–13^, only a few recent methods^14–17^ capture dynamic regulatory relationships (i.e., transcription factor – target gene; TF-TG) by ordering cells along pseudotime trajectories. Although effective in capturing causal TF-TG interactions when pseudotime is accurately inferred^16^, the biological complexity poses several fundamental challenges to their accuracy and interpretability. Pseudo-temporal ordering complicates analysis of diverse populations because it assumes temporal variation reflects time-delayed regulation within individual cells, an assumption that may not hold in heterogeneous systems. Moreover, TF-TG rewiring along trajectories further complicates interpretation, as connections may be cell-type-or stage-specific rather than active throughout development.

To address this, we combined static GRNs inferred for neurogenic lineages with their pseudotime to evaluate the temporal dynamics of biologically interpretable regulatory subnetworks comprising regulons and gene modules. By focusing on these subnetworks, we not only reduce the computational complexity of tracking individual TF– TG rewiring events but also capture combinatorial gene regulatory effects as coherent biological functional units^18^. Furthermore, we introduced a dynamic score metric as a quantitative measure of how regulatory subnetwork activity along the developmental trajectory, allowing us to prioritize subnetworks most relevant to neurogenesis (**Table S1**). We present an extensive resource for early mammalian neurogenesis, emphasizing dynamic regulons and gene co-regulatory modules. This resource includes 18 human neurogenesis GRNs spanning three regional hierarchies and collectively representing the entire human brain, along with 4 non-human GRNs to facilitate cross-species comparisons. In addition to characterizing dynamic regulons and co-regulatory gene modules across neurogenic lineages, we investigated the overrepresentation of specific regulons within risk genes associated with over 30 neurological disorders and developmental traits. This analysis establishes connections between neurogenic lineages, their associated regulons, and disease vulnerability. An accompanying online platform (https://daifengwanglab.shinyapps.io/devGRNDB/) is provided for interactive exploration of these networks.

## RESULTS

### Identification and characterization of the dynamics of regulatory networks in the developing mammalian brain

Delineating the dynamics of neurodevelopmental regulatory networks presents several fundamental computational challenges, primarily due to overlapping differentiation and maturation processes that confound cell-state resolution. This complexity necessitates a robust and quantifiable metric to assess the degree of association between regulatory features and neurodevelopment. To address this, we developed a computational framework (**Figure 1A**) for identifying gene regulatory subnetworks—including regulons (TFs and their TGs) and gene modules—that exhibit dynamic variation across biological processes.

**Figure 1.**
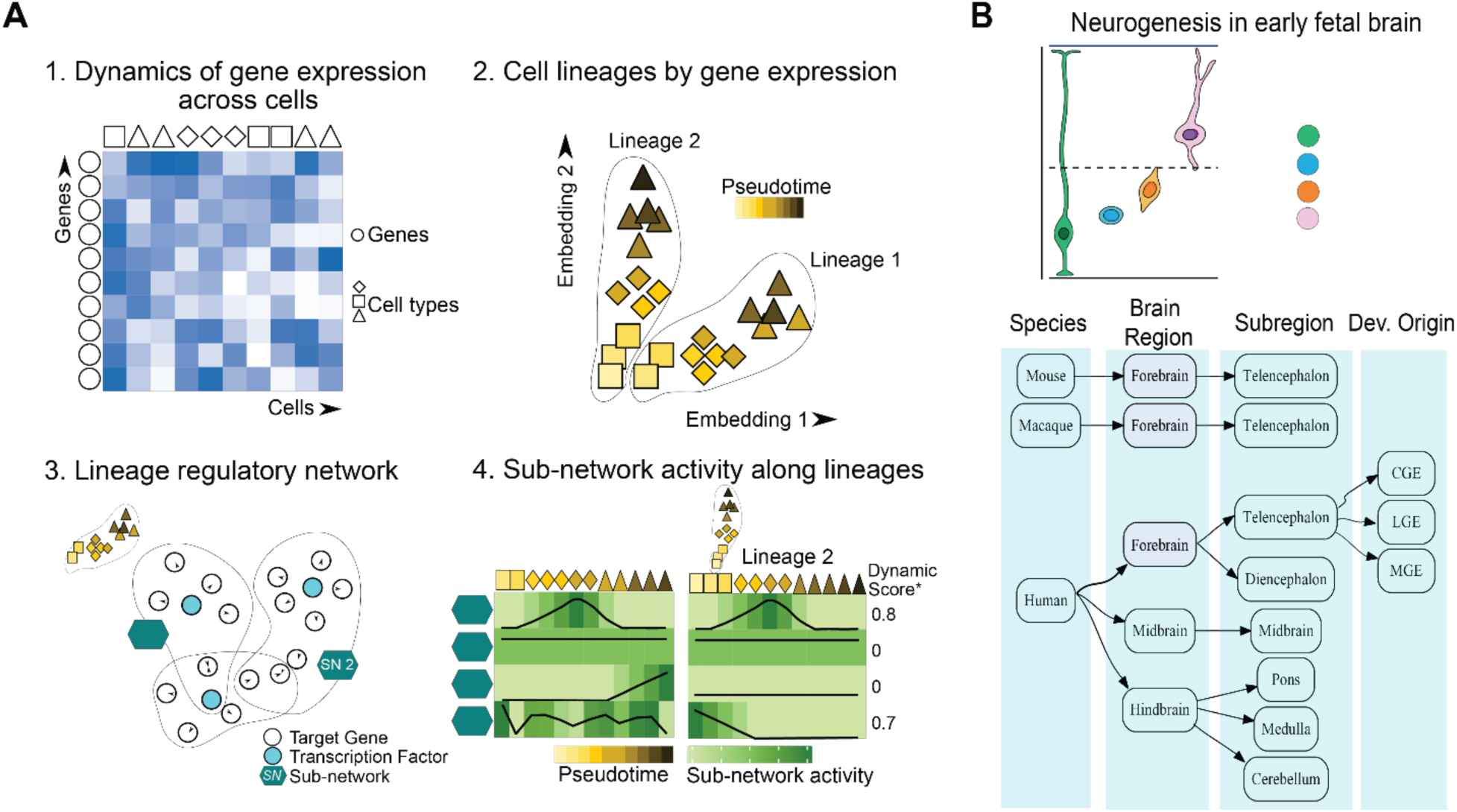
Constructing a dynamic gene regulatory network resource using the developmental single cell data. **A.** Computational workflow for dissecting dynamics of gene regulation in neurogenesis 1. Input related single-cell gene expression data. 2. Lineage assignments via cell fate probabilities and pseudotime inference of single cells. 3. Construction of gene regulatory networks for the lineage and identification of regulatory subnetworks with potential expression changes across lineage. Regulatory subnetworks could either be regulons (i.e., transcription factor and its target genes) or derived gene modules. 4. Quantification and prioritization of dynamic regulatory subnetworks for lineages. **B.** Three-tiered resource integrates whole-brain and cross-species datasets to identify dynamically regulated subnetworks throughout early human neurodevelopment. Human lineages include 3 broad regions, 6 subregions, each with glutamatergic and GABAergic lineages, and three GABAergic subtype lineages. Species include human, macaque, and mouse. IPC, intermediate progenitor cell; CGE, caudal ganglionic eminence; LGE, lateral ganglionic eminence; MGE, medial ganglionic eminence.

This framework consists of four main steps (**Figure 1A**). We begin with single-cell gene expression data, along with accompanying metadata, such as cell-type annotations and temporal information. Second, we infer cellular trajectories, or pseudotime, to model the progression of cellular states (e.g., developmental periods) across the given biological process. In cases with multiple lineages (e.g., glutamatergic or GABAergic neurogenesis), we apply lineage inference tools to estimate fate probabilities prior to pseudotime estimation, enabling lineage-specific trajectory inference. Third, we infer GRNs independently for each lineage, allowing extraction of lineage-specific regulatory subnetworks (**Figure 1A, step 3**). Fourth, we quantify subnetwork activity for each cell via enrichment of gene features incorporated within each subnetwork. This reveals both the relative activity of distinct subnetworks within individual cells and how the behavior of each subnetwork (i.e., regulon or module) varies across cellular landscapes (**Figure 1A, step 4; Methods**). Subnetwork activity may follow simple trends (e.g., monotonic increase or decrease), complex patterns (e.g., biphasic), or appear non-dynamic (i.e., constant or random). Given this variability, standard correlation-based association metrics are insufficient. Therefore, we introduce a dynamic score by adopting Moran’s I statistic—a spatial autocorrelation metric—to measure how similarly valued subnetwork activities are distributed across pseudo-temporally ordered cells. Higher scores indicate pronounced, non-random variations, reflecting stronger temporal associations likely contributing to transcriptional programs relevant to neurogenic lineages.

Using this framework, we conducted a comprehensive analysis of human brain development during the first trimester (PCD 35-98)^19^, together with developmentally matched stages in macaque (PCD 37-78)^20^ and mouse (PCD 7-18)^21^, thereby uncovering regulatory dynamics across brain regional hierarchies and species (**Figure 1B, Figure S1**).

### Developmental gradients and regionalization of human neurogenic regulatory networks

We first evaluated whether our methodology could identify the principal regulons driving transcriptional programs for neuronal differentiation and broad regional identity establishment in the developing human brain. Our analysis identified three major neurogenic lineages corresponding to the forebrain, midbrain, and hindbrain (**Figures 2A-C**). Each lineage displayed a pseudotime continuum of cell states from radial glial cells (RGs) to intermediate progenitor cells (IPCs), early neuroblasts, and neurons, consistent with the progression of neuronal differentiation (**Figures 2A, B**).

**Figure 2:**
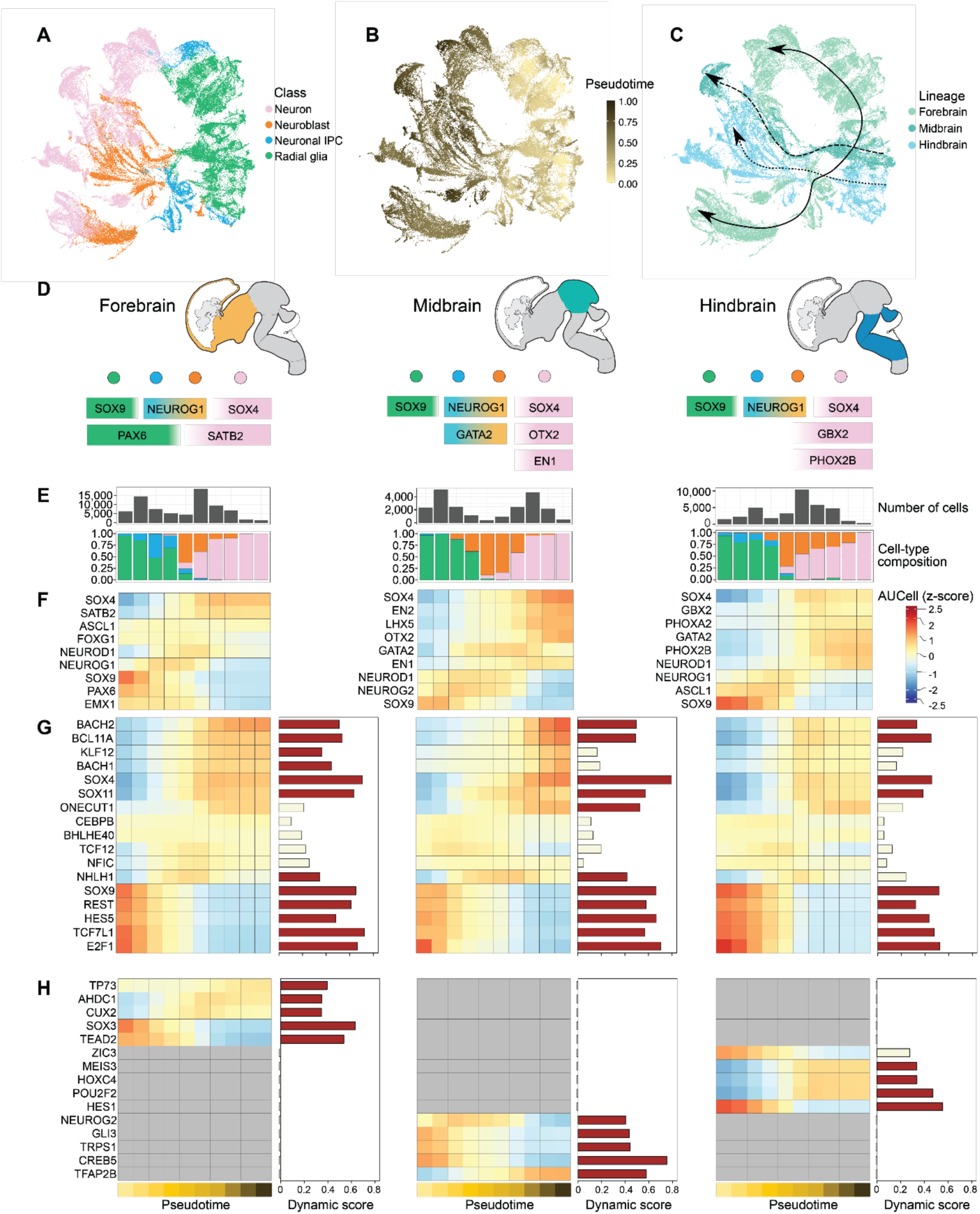
**Developmental gradients and regionalization of dynamic regulatory subnetworks in early human neurogenesis. A–C**. UMAP visualizations of pseudobulk cells spanning the first trimester human brain, colored by cell type (**A**), pseudotime (**B**), and lineage assignment (**C**). Arrows indicate neurogenic trajectories identified for the forebrain (solid line), midbrain (dashed line), and hindbrain (dotted line). **D.** Schematic of key transcription factors essential for neuronal differentiation and regional identity specification in the forebrain (left), midbrain (center), and hindbrain (right) during early brain development. Colors represent cell type specific expression along neurogenesis. Green, radial glia; blue, neuronal IPC; orange, neuroblast; pink, neuron. **E.** Bar plots depicting the number of cells (top) and cell type composition colored by cell type (bottom) across pseudotime bins for each brain. **F.** Pseudotemporal variation in regulon activity for key transcription factors shown in **Figure 2D**. **G–H.** Pseudotemporal profiles of regulon activity for selected highly dynamic, lineage-conserved transcription factors (**G**), and lineage-specific transcription factors (**H**). Bar plots denote dynamic scores, with red indicating values greater than 0.3 and tan indicating values less than 0.3.

Our approach uncovered gradients of regulon activity encompassing well-characterized TFs essential for both general neuronal differentiation and the establishment of regional identities within the forebrain, midbrain, and hindbrain during early brain development (**Figures 2D-F**). Throughout neurogenesis, TFs regulate the balance between neural stem cell self-renewal, progenitor expansion, and neuronal differentiation^22^. We identified the PAX6 regulon—a master forebrain regulator that promotes neural stem cell proliferation and neurogenesis^23^—and the SOX9 regulon, crucial for the maintenance and proliferation of neural progenitors^24^, as highly enriched in forebrain RGs, with their activity progressively declining across pseudotime (**Figure 2F**). Additionally, we found the NEUROG1 regulon to be highly enriched in IPCs, consistent with its established role as a PAX6-activated proneural basic helix-loop-helix (bHLH) TF promoting neuronal differentiation^25^ (**Figure 2F**). Furthermore, enrichment of the SOX4 regulon in postmitotic neurons aligned with its downstream role in terminal neuronal differentiation^26^ (**Figure 2F**).

In addition to regulons associated with general neuronal differentiation, we identified well-characterized region-specific regulons that define early brain regional identity. These include the SATB2 regulon, a chromatin remodeler and marker of developing cortical excitatory neurons^27^, and the OTX2 regulon, which defines the anterior neural plate and is required for the regionalization and patterning of the midbrain during early embryogenesis^28^ (**Figures 2D-F**). The EN1 and EN2 regulons essential for midbrain neuron development and survival were also found to be highly enriched in midbrain neurons^29^ (**Figures 2D-F**). In the hindbrain, we observed enrichment of the GBX2 regulon, necessary for the specification and formation of the anterior hindbrain (rhombomeres 1-3)^30^, and the PHOX2B regulon, which is critical for autonomic and visceral motor neuron development^31^ (**Figures 2D-F**). Together, these results validate the sensitivity of our approach in detecting major neurogenic regulons and demonstrate the capacity to identify both broad-acting and region-restricted regulons fundamental to human brain development.

### Gene regulatory network dynamics of neurogenic transcription factors across human development

In addition to confirming established neurogenesis regulators, we uncovered novel and putative drivers of neuronal differentiation conserved across all three neurogenic lineages via dynamic score metrics that reflect regulon activity changes across differentiation and maturation (**Methods**).

To evaluate which modality—transcription factor gene expression or regulon activity—more accurately captured the pattern of transcriptional regulation during neuronal differentiation, we directly compared their dynamics across pseudotime. As expected, both exhibited broadly concordant temporal variation (**Figures S2A-B**). However, the magnitude of dynamic scores was significantly greater for regulon activity than for TF gene expression (Wilcoxon signed rank test, p ≤ 2.2e-16, **Figure S2C**). This discrepancy likely arises from the inherent sparsity and stochastic dropouts associated with single-cell RNA-sequencing, which compromises the reliability of single-gene measurements^32^. In contrast, regulon activity aggregates expression across downstream targets, yielding a more stable and biologically interpretable signal that is less sensitive to technical noise. Notably, the reduced dynamic scores derived from gene expression were not uniformly observed across all regulons, indicating that the extent of this attenuation is context-dependent and likely influenced by individual regulon dynamics and TG composition (**Figure S2C**). Furthermore, to test the robustness of our framework, we performed a sensitivity analysis using alternative GRN inference methods, revealing varying degrees of regulon overlap (**Figure S3A**), yet a strong concordance of regulon dynamic scores (**Figures S3B-C**). This robustness likely stems from our reliance on aggregated regulon activity metrics rather than individual network TF-TG links, which preserves core regulatory patterns despite methodological differences in network construction. Together, these findings underscore the advantage of regulon-based approaches for resolving cell state transitions, particularly in dynamic processes such as neuronal differentiation, when transcriptional programs are driven by coordinated regulatory mechanisms rather than discrete changes in the expression of single genes.

We then identified several lineage-conserved and highly dynamic regulons implicated in RG proliferation and cell cycle regulation, including HES5, TCF7L1, and E2F1, which were significantly enriched in RGs and IPCs (**Figure 2G**). In addition, we found regulons enriched in IPCs and immature neuroblasts, including many proneural bHLH TFs (BHLHE40, TCF12, NHLH1), known to regulate early neuronal differentiation programs^33^ (**Figure 2G**). While prior studies have primarily emphasized Nuclear Factor 1 (NFI) isoforms, such as NFIA and NFIB, for their roles in neuronal differentiation^34^, our analysis uncovered NFIC regulon enrichment within forebrain IPCs and immature neuroblasts (**Figure 2G**), suggesting functional redundancy among NFI isoforms in immature neurons and implicating NFIC as an understudied factor in human neurogenesis.

Notably, IPC-and immature neuroblast-enriched regulons exhibited broader, less defined gradients compared to those associated with cell cycle and proliferation (**Figure 2G**), suggesting a gradual onset and offset of neuronal transcriptional programs, in contrast to the abrupt on-off dynamics characteristic of proliferation and cell cycle regulons. This aligns with a gradual increase in neuronal gene expression during early differentiation, followed by further refinement through highly dynamic terminal neuronal differentiation regulons, such as SOX4, SOX11, and BACH1 (**Figure 2G**).

Next, we identified the top 10 highly dynamic regulons (via dynamic regulon scores) conserved across all three neurogenic lineages (**Figure S2D**), recapitulating core regulators of neural proliferation (e.g., HES5, E2F3, HMGA2) and differentiation (e.g., SOX4) along with highlighting GATA3 as both highly dynamic and enriched in neurons across all broad regions (**Figure S2D**). Although previously linked to specific neuronal subtypes^35^, our findings suggest a broader, conserved role for GATA3 in driving neuronal differentiation throughout the CNS.

Following the identification of known and candidate regulators of general neurogenic processes, we sought to define region-enriched regulons by identifying the top five lineage-specific regulons based on their dynamic scores (**Figure 2H**). This revealed several well-characterized regional transcriptional regulators, including CUX2, which is enriched in developing upper-layer cortical excitatory neurons^36^; GLI3, essential for the coordination of anteroposterior and dorsal-ventral patterning of the midbrain through integration of Sonic hedgehog (SHH) and FGF8 signaling^37^; and MEIS3 and HOX4, key drivers in hindbrain induction and anteroposterior rhombomere specification^38–40^. A summary of all the broad region regulons is provided in **Table S7**.

We then constructed a bipartite network connecting highly dynamic regulons (dynamic score > 0.4) to their respective lineages **(Figure S2E**). This network revealed both conserved (e.g., E2F family TFs) and region-specific regulons, including SATB2 and PAX6 in the forebrain, GSX1 and TFAP2B in the midbrain, and HOXA3 in the hindbrain, demonstrating spatially resolved dynamic regulatory programs across brain development (**Figure S2E**). Beyond known regulators, we identified candidate region-enriched regulons with limited prior association to brain regionalization: TEAD2 in forebrain progenitors, CREB5 in midbrain, and ZIC3 in hindbrain, suggesting yet uncharacterized roles for these factors in specifying neuron regional identity (**Figure 2H, Figure S2E**).

Interestingly, several top forebrain neuron-enriched regulons identified in our analysis have been previously implicated in neurodevelopmental disorders (NDD), underscoring the potential clinical significance of our gene regulatory resource. Among these is CUX2, a highly dynamic regulon enriched in the forebrain, with pathogenic mutations known to underlie a neurodevelopmental syndrome characterized by drug-resistant epilepsy and intellectual disability^41^ (**Figure 2H**). Similarly, TP73 was identified as a dynamic forebrain-enriched regulon, with loss-of-function variants established contributions to multiple developmental forebrain malformations, including lissencephaly, hippocampal dysgenesis, and white matter abnormalities^42^ (**Figure 2H**).

Among the most dynamic regulons in the forebrain was AHDC1, a master regulator of chromatin organization and gene expression^43^ **(Figure 2H**). Pathogenic variants in *AHDC1* are causative of Xia-Gibbs syndrome, a multisystem neurodevelopmental disorder characterized by callosal hypoplasia, delayed myelination, intellectual disability, and developmental delay^44^. Despite well-established clinical associations, AHDC1’s role in early brain development remains poorly understood. This is largely attributable to the limited availability of functional models, as Ahdc1 knockout mice exhibit early embryonic lethality^45^. To further dissect the AHDC1 regulatory network in early human brain development, we constructed an AHDC1 GRN subnetwork by isolating upstream regulators and downstream target genes in the developing forebrain **(Figure S2F)**. These results not only substantiate a critical connection between AHDC1 and regional neuronal differentiation but also provide a gene regulatory framework for dissecting the molecular etiology of AHDC1-associated neurodevelopmental disorders. Notably, AHDC1 targets identified during this pivotal developmental window include *NR2F2*, *KLF7*, *FCGR1A*, enabling the prioritization of candidate effectors that may contribute to the structural and functional neurological deficits in Xia-Gibbs syndrome (**Supplementary Figure 2F**).

### Dynamic gene modules of brain regionalization

Our regulon-based analysis presents potential avenues for further study or experimental manipulation of key transcriptional regulators. However, identifying a dynamic regulon alone does not inherently convey its functional significance beyond a broad neurogenic association. For instance, a highly dynamic but poorly characterized regulon may emerge without a clear biological relevance. To address this, we extended our approach to identify dynamic gene modules, defined as co-regulatory network modules that cluster genes based on shared regulatory patterns, revealing coordinated biological processes along individual neurogenic lineages (**Methods; Table S5**).

As expected, many dynamic modules showed overlapping functional annotations among the forebrain, midbrain, and hindbrain (**Figures S4A-C**), encompassing fundamental biological processes, including translation, cell cycle progression, and post-translational protein modifications, as well as processes specifically related to neuronal differentiation and maturation, such as neuron projection morphogenesis and synapse development (**Figures S4A-C**).

Interestingly, we found midbrain and hindbrain modules with significant functional enrichment for WNT signaling (**Figures S4B, C**), consistent with the well-characterized anteroposterior (AP) gradient of WNT signaling during vertebrate brain development, with low WNT activity in anterior (forebrain) and progressively higher levels toward more posterior (hindbrain) regions^46^. To further assess overlap and region-specificity among WNT signaling-associated gene modules, we constructed a differential GRN, highlighting conserved and region-enriched WNT signaling-associated genes for the midbrain (Module M3) and hindbrain (Module M9) (**Figure S4D**). This demonstrates the strength of our dynamic module-based approach in resolving both conserved and regionally specialized morphogenetic processes that underpin neurodevelopmental regionalization. Collectively, this dynamic gene module analysis provides a valuable resource for the neuroscience community, offering new insights into the conserved and region-specific gene programs that coordinate brain patterning and development.

### Region and lineage-specific dynamic regulatory subnetworks in the developing human hindbrain

To further dissect the regulatory networks underlying hindbrain regional specification, we subset hindbrain cells and identified neurogenic lineages corresponding to the developing cerebellum, pons, and medulla (**Figure S1**). Importantly, at this regional hierarchy, we resolved two distinct lineages within each hindbrain subregion, corresponding to glutamatergic and GABAergic neurogenic lineages (**Figure 3A, Figure S5A, and Figure S1, Table S7**). In the glutamatergic lineages, we identified a cohort of dynamic regulons conserved across all three hindbrain regions (**Figure 3B**). Many of these conserved regulons, including SOX9, E2F2, and HMGA2, are involved in cell cycle regulation and neural progenitor proliferation, underscoring the importance of tightly controlled proliferative mechanisms during early neuronal lineage cell expansion.

**Figure 3.**
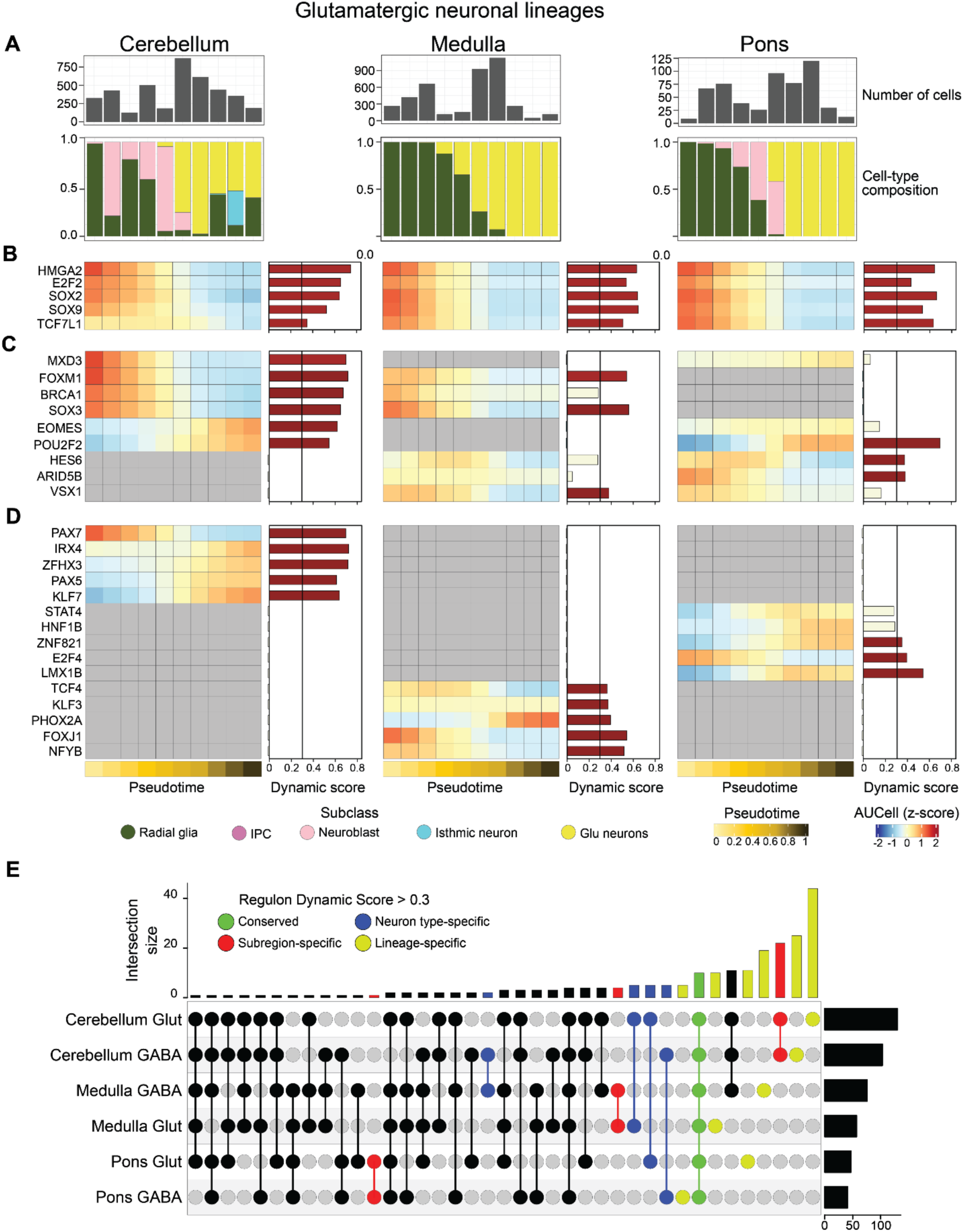
Regulon dynamics across human hindbrain subregion neurogenic lineages. **A.** Bar plots depicting the number of cells (top) and cell type composition colored by cell type (bottom) across pseudotime bins for the cerebellum, medulla, and pons lineages. RG, radial glia; IPC, intermediate progenitor cells; Glu neurons, glutamatergic neurons. **B–D**. Pseudotemporal profiles of regulon activity z-scores of selected regulons classified as: conserved in all three lineages and highly dynamic (**B**), pairwise conserved (i.e., present in two of the three lineages) (**C**), and unique to individual lineages (**D**). Bar colors indicate dynamic score, with red denoting scores > 0.3 and tan < 0.3. **E.** UpSet plot illustrating overlaps among dynamic regulons (score > 0.3). Green marks regulons conserved across all six lineages; red highlights those specific to hindbrain subregions regardless of neuronal identity, blue indicates regulons restricted to a neuronal type (i.e., GABAergic neurons), and yellow indicates regulons specific to each lineage.

Next, we characterized region-enriched dynamic regulon patterns by pairwise comparisons and region-specific regulons (**Figures 3C, D**). This revealed cerebellar-specific enrichment of the PAX7 regulon (**Figures 3D**), a TF expressed in the alar plate of rhombomere 1^47^, a segment of the developing hindbrain comprising the cerebellar primordium. Additionally, the medulla demonstrated selective enrichment of the PHOX2A regulon (**Figure 3D**), an essential TF for the development of autonomic and visceral reflex circuit neurons in this region, with PHOX2A deficiency causing impaired development and disrupted connectivity of medullary neuron populations^31,48^. Furthermore, the LMX1B regulon was found to be enriched in the pons (**Figure 3D**). LMX1B, a LIM homeodomain TF, is expressed in the pontine nuclei and is essential for the development of pontine serotonergic neurons^49^. Analysis of GABAergic lineages similarly identified conserved, pairwise, and region-specific regulons across the developing hindbrain subregions, highlighting known (i.e., HOXA6 in medulla) and candidate (i.e., HIVEP2 in cerebellum) regulators of GABAergic neurogenesis (**Figures S5B-D**).

While region-enriched regulons help specify discrete neuronal populations, the broader architecture of dynamic GRNs frequently relies on distributed, combinatorial control through interactions among multiple TFs^50^. Rather than the rewiring of a single regulator, shifts in lineage identity and regional specification are, in general, mediated by coordinated regulation across numerous network nodes, reflecting the inherent complexity of developmental GRNs^51^. Furthermore, our identification of both glutamatergic and GABAergic lineages within each hindbrain subregion enabled us to investigate whether brain region or neuronal lineage (i.e., GABAergic versus glutamatergic lineages) primarily drives conservation or divergence in regulon composition. To address this, we systematically evaluated all highly dynamic regulons (dynamic score >0.3) across the six hindbrain lineages, quantifying the number of conserved and lineage-specific regulons (**Figure 3E, Figure S5E**).

We found a predominance of lineage-specific dynamic regulons relative to subregion-conserved regulons, highlighting the principal connection of individual lineage identity to regulatory landscapes (**Figure 3E**). Notably, some subregion-conserved dynamic regulons, such as those common to the medulla, were more prevalent than neurotransmitter phenotype-shared dynamic regulons, such as the regulons common to all GABAergic neurons irrespective of region (**Figure 3E**). This suggests a complex regulatory architecture characterized by a high degree of lineage specificity governing neuronal subtype specification, alongside distinct regulons that define hindbrain subregional identity.

The cerebellum exhibited the largest number of subregion-conserved and lineage-unique dynamic regulons, encompassing both GABAergic and glutamatergic lineages (**Figure 3E**). This striking enrichment underscores the cerebellum’s unique and complex regulatory landscape during neurodevelopment, consistent with distinct transcriptomic profiles in the adult cerebellum^52^. Furthermore, we identified dynamic regulons conserved across all six lineages, representing a core set of regulons fundamental to proliferation and general neuronal development (**Figure 3E and Figure S5F**). Our analysis revealed a comparatively smaller set of dynamic regulons dedicated to the specification of neurotransmitter phenotypes, demarcating GABAergic versus glutamatergic cell fates (**Figure 3E**). Collectively, these findings emphasize that while major neurotransmitter systems are utilized across diverse brain regions - e.g., pontine versus cerebellar GABAergic neurons - the underlying dynamic regulons that specify the connectivity and functional properties of these neurons diverge substantially, reflecting regional context-dependent regulatory mechanisms.

To further elucidate the key regulons underlying neuronal lineage and regional specialization, we constructed a bipartite network to visualize the relationship between highly dynamic regulons and hindbrain lineages (**Figure S5E**). This identified MYCN as a cerebellar glutamatergic lineage-specific regulon (**Figure S5E**). MYCN is known to play a critical role in cerebellar development by regulating the proliferation and maturation of granule cell progenitors (GCPs), with functional loss resulting in reduced GCP proliferation and cerebellar hypoplasia^53^. Furthermore, the dynamic regulon RXRG, a nuclear receptor that mediates retinoic acid signaling, was significantly enriched in both glutamatergic and GABAergic lineages of the medulla (**Figure S5E**). This finding aligns with the established role of retinoic acid in rostrocaudal hindbrain patterning, yet also highlights the comparatively underexplored role of RXRG relative to other retinoic acid receptors implicated in hindbrain development. Collectively, these findings delineate the regulatory programs that contribute to the specification and maturation of distinct neuronal populations within the developing hindbrain.

### Divergent gene regulatory subnetworks shape human telencephalic GABAergic neurogenic processes and associate with neurodevelopmental disorder genetic risk

While our computational framework defined the principal glutamatergic and GABAergic lineages within hindbrain subregions, the resolution of these lineages remained relatively coarse. This is primarily due to the molecular and cellular heterogeneity inherent to the developing brainstem, which gives rise to a diverse array of neuronal subtypes organized into functionally specialized nuclei with complex and highly divergent patterns of connectivity^54^. Therefore, we performed a more granular characterization of select neuronal lineages, identifying discrete, molecularly defined neuron subtypes at a finer hierarchical resolution (**Figure 1B**). The ganglionic eminences (GE) - transient proliferative structures of the ventral telencephalon - serve as the origin for the majority of human telencephalic GABAergic neurons^55^. These include the medial (M), caudal (C), and lateral (L) GEs, as well as the developing ventromedial forebrain (VMF), each generating distinct GABAergic neuron subclasses with unique developmental trajectories, molecular identities, and functional properties.

Focusing on telencephalic GABAergic lineage cells, we annotated neuroepithelial stem cells (NESC), radial glia (RG), intermediate progenitor cells (IPC), and GABAergic neurons of the telencephalon and performed lineage inference, resolving major developmental trajectories emerging from the GEs. We identified distinct trajectories corresponding to MGE, LGE, and CGE lineages **(Figures S6A, B**), marked by *NKX2-1* and *LHX6* in MGE-derived, *MEIS2* and *ISL1* in LGE-derived, and *PROX1* and *SP8* in CGE-derived neurons (**Figure S6A**). All lineages displayed robust expressions of *DLX1* and *DLX2*, established markers of maturing and migrating GABAergic interneurons^56^ (**Figure S6A**). VMF neurons were excluded from the analysis due to their limited availability, which precluded reliable trajectory reconstruction. At this refined level of lineage resolution, we conducted a systematic analysis of dynamic regulatory subnetworks governing telencephalic GABAergic neuron subtype development.

We employed regulon-and gene module-based analyses to delineate functional similarities and differences across subtype-specific neuronal GRNs (**Figure 4A and Figures S6B-E**). However, this presented an inherent computational challenge in deriving consensus gene modules representative of all three GRNs (i.e., clustering of a multi-layer network, each layer representing a neuronal subtype). To overcome this, we leveraged the hierarchical regional architecture in our dataset (**Figure 1B**) and posited that the telencephalic GABAergic lineage, defined at the subclass level, offers a biologically grounded consensus framework encompassing the three GABAergic subtype lineages (**Figure 2A**). This enabled us to trace gene module dynamics along a nested regulatory topology, linking molecular patterns to functional annotations and associations with neurodevelopmental disorders.

**Figure 4.**
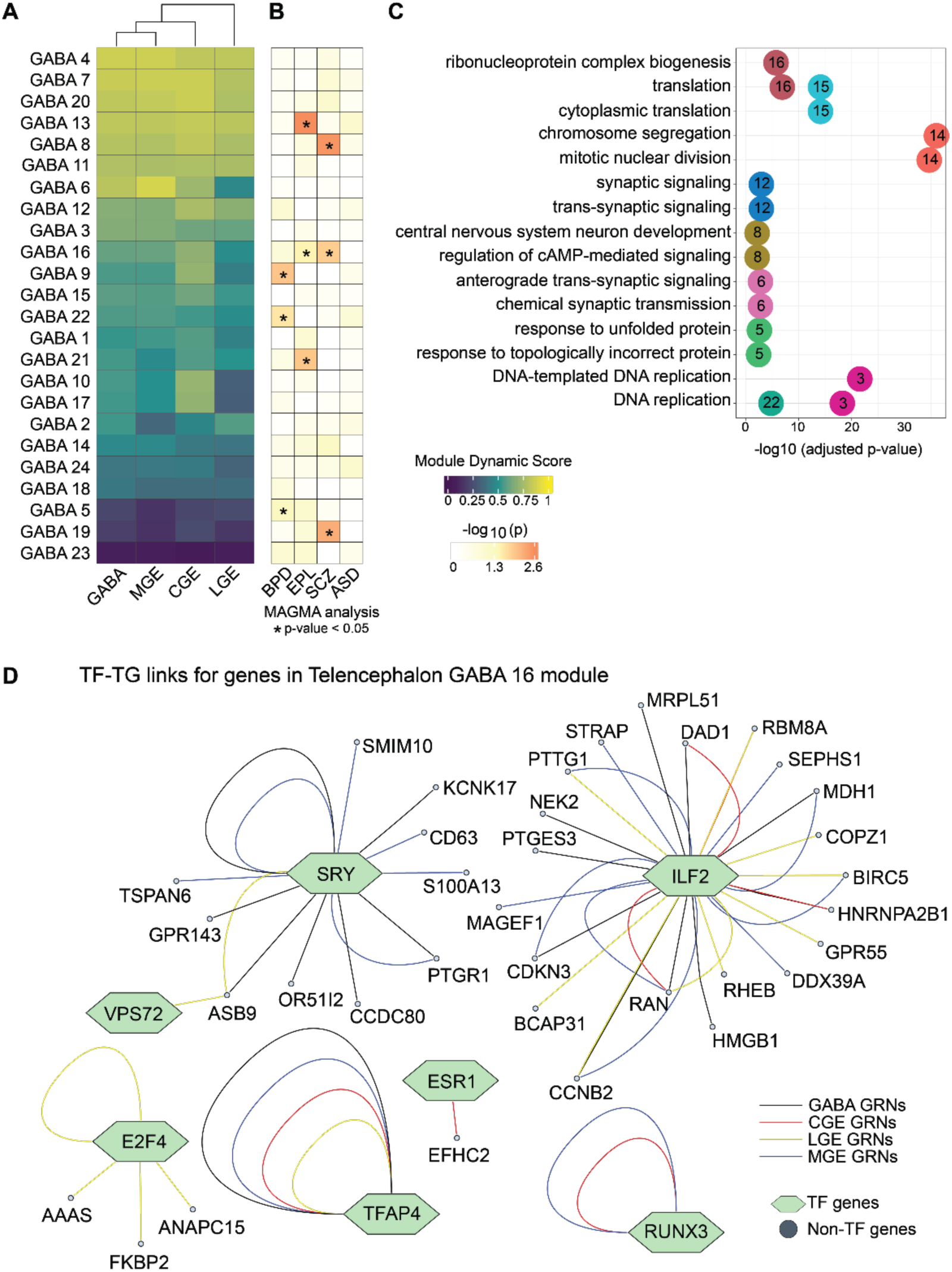
Linking telencephalic GABAergic neurogenesis to neurodevelopmental disorder risk. **A.** Heatmap of module dynamic scores for co-regulatory gene modules (e.g., GABA-1, GABA-2) derived from GABAergic neurogenesis. Module activity was subsequently estimated across subtype lineages (CGE, LGE, MGE), and their module dynamic scores were calculated. GABA, GABAergic neuron subclass level lineage; CGE, caudal ganglionic eminence lineage; MGE, medial ganglionic eminence lineage; LGE, lateral ganglionic eminence lineage. **B.** Heatmap showing MAGMA-derived disease enrichment p-values for these modules, across four neurodevelopmental disorders: ASD (autism spectrum disorder), BPD (bipolar disorder), EPL (epilepsy), and SCZ (schizophrenia). **C.** Gene Ontology (GO: Biological Process) terms enriched in each co-regulatory module. **D.** Network visualization of GABA-16 module genes across the four neurogenic lineages. Transcription factors are shown as hexagons; non-transcription factors as circles. Edge colors reflect lineage-specific transcription factor–target associations.

We therefore constructed a GABAergic neuron co-regulatory network and applied hierarchical clustering to identify gene modules (**Figure 4A, Methods**). Dynamic scoring of gene modules across GABAergic lineages revealed differential dynamics of consensus modules (**Figure 4A**). Given the role of telencephalic GABAergic neurons in NDDs, we assessed disease associations via linkage disequilibrium (LD)-aware enrichment analysis of genome-wide association study (GWAS) traits^57^. This identified significant associations of developmental GABAergic neuron gene modules with GWAS risk loci for schizophrenia (SCZ), bipolar disorder (BPD), and epilepsy (EPL), whereas no significant enrichment was observed for autism spectrum disorder (ASD) (**Figure 4B)**. Complementarily, we performed GO enrichment analysis to link gene modules to biological processes (**Figure 4C**). Together, this approach facilitated the characterization of highly dynamic gene modules within GABAergic neuronal lineages and their association with neurodevelopmental disorder risk (**Figures 4A-C)**.

A particularly illustrative example of this approach is the GABA-8 module, which exhibited pronounced module activity across all developmental lineages, was enriched for neurodevelopmental processes and cAMP-mediated signaling pathways, and was significantly associated with schizophrenia risk loci (**Figures 4A-C**). These findings align with the established role of cortical GABAergic interneuron abnormalities in the pathophysiology of schizophrenia, particularly in cortical microcircuits of the frontal cortex^58^. Dysregulation of cAMP signaling - a critical modulator of synaptic function and receptor expression - has also been documented in the frontal cortex of individuals with schizophrenia^59^. Together, our findings identify a highly dynamic gene module involved in GABAergic neuron development and cAMP-mediated signal transduction (GABA-8 module) that is tightly linked to genetic risk for schizophrenia (**Figures 4A-C**). These results identify convergent neurodevelopmental mechanisms linking GABAergic neuron maturation and intracellular signaling to developmental cortical circuit assembly and schizophrenia susceptibility.

In addition to uncovering gene modules that were highly dynamic across all developmental lineages, our analyses further revealed modules with lineage-specific dynamic profiles. For example, the GABA-6 module was highly dynamic in the CGE and MGE lineages, but with markedly reduced activity in the LGE lineage (**Figure 4A**). Functional annotation indicated enrichment for synaptic processes and anterograde trans-synaptic signaling, underscoring lineage-dependent maturation and synaptic specialization among telencephalic GABAergic neurons (**Figure 4C**).

We also identified gene modules (GABA-5, GABA-9, GABA-22) that were relatively less dynamic along pseudotime yet demonstrated enrichment for bipolar disorder risk loci (**Figures 4A, B**). These modules highlight critical cellular processes involved in genomic stability and proteostasis during neurogenesis, suggesting that disruptions in early DNA replication fidelity and proteostatic regulation may contribute to bipolar disorder vulnerability (**Figure 4C**). Identification of BPD-associated genetic risk within these functional pathways supports the hypothesis that early developmental disturbances influence neural circuit formation and resilience, conferring developmental differential susceptibility to BPD.

Furthermore, GABA-16 emerged as a crucial gene module due to its pronounced activity and significant enrichment for risk loci associated with both schizophrenia and epilepsy (**Figures 4A, B**). Functionally, GABA-16 was enriched for ribonucleoprotein complex biogenesis and translation **(Figure 4C**), aligning with prior studies on the temporal regulation of translation during neurogenesis^60^.

To elucidate the regulatory architecture within the GABA-16 gene module, we identified TF-TG link specificities across GABAergic lineages (**Figure 4D**). The TF and RNA-binding protein ILF2 emerged as a central regulatory hub, exhibiting both shared and divergent connectivity patterns across GABAergic lineages (**Figure 4D**). These findings underscore the essential role of translation in the development and integration of telencephalic GABAergic neurons, and link dynamic neurodevelopmental gene programs of translational control to schizophrenia and epilepsy genetic risk.

Together, these results demonstrate the applicability of our gene regulatory subnetwork resource to resolve the regulatory architecture of human telencephalic GABAergic neuron development. This resource facilitates the linkage of developmental regulatory subnetworks to cellular functions and neurodevelopmental disorder genetic risk, providing insights into the molecular pathways underpinning cortical circuit assembly and disease etiology.

### Mapping neurological disorder risk genes to dynamically varying regulatory subnetworks

Building on the GABAergic lineage modules and their roles in neurological disorders, we next sought to identify which neurogenic regulators are most affected or potentially drive neurological disorders. Therefore, we extended our analysis to examine lineage associations with over 30 neurological disorders, cortical malformations, and developmental traits. Using a hypergeometric test, we identified highly dynamic regulons significantly overrepresented in curated disease gene sets (**Methods, Table S6**). These lineage–regulon–disorder associations (**Figure 5A)** highlight neurogenic regulons potentially disrupted in neurological disease and reveal potential region-based signatures. Among the most enriched were SOX11, SOX4, NFIB, and BCL11A, previously linked to intellectual disability, facial anomalies, microcephaly (SOX4^61^, SOX11^62^), and ASD (BCL11A^63^).

**Figure 5:**
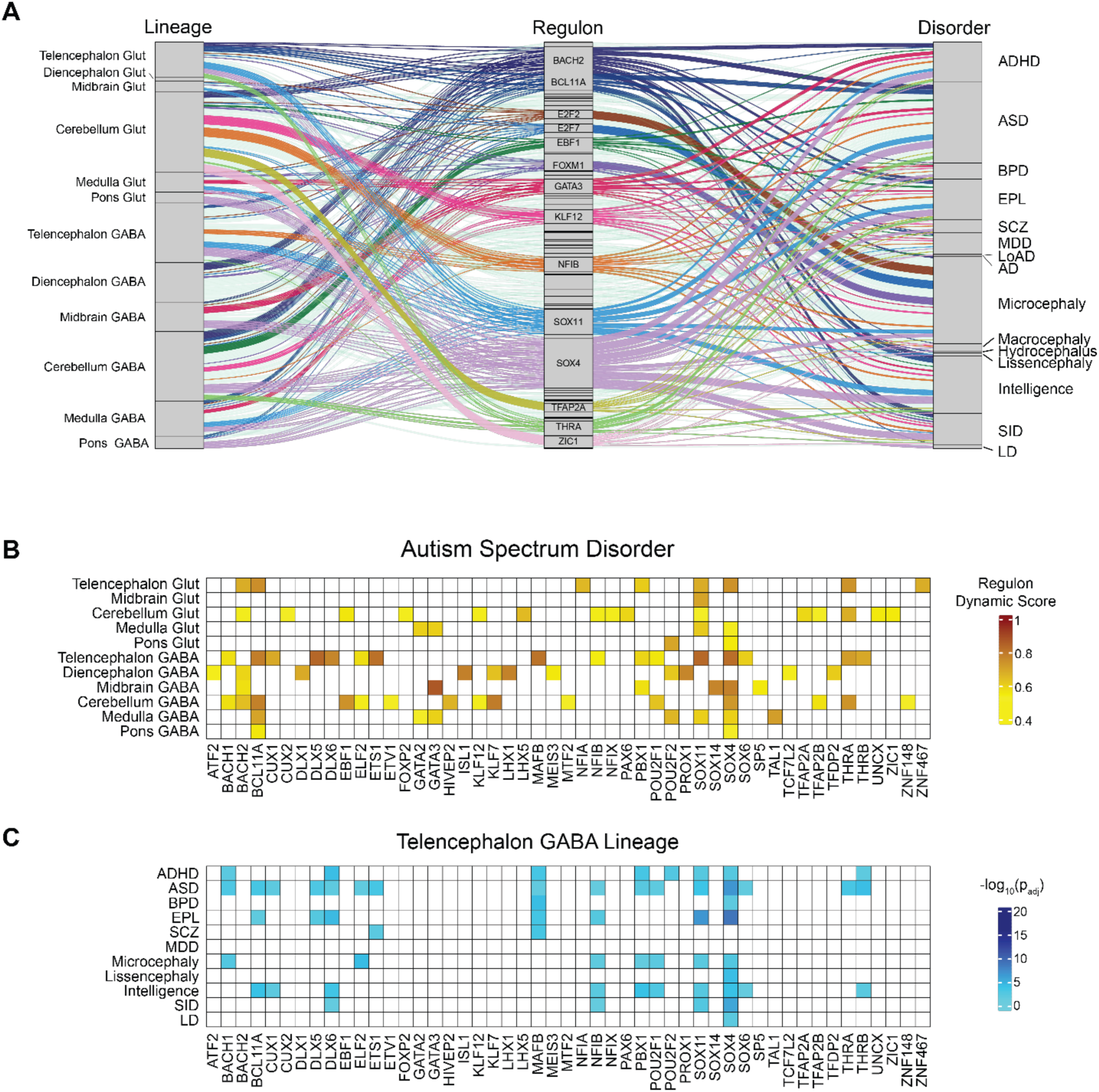
L**i**neage**-Disease risk gene association. A.** A Sankey plot depicting the associations connecting lineages, regulons, and disorders. Lineage-to-regulon associations were made using the regulon dynamic scores, and regulon-to-disorders associations were identified through the overrepresentation of regulons on disorder risk genes via a hypergeometric test. All the links that have an adjusted p-value less than 0.05 and a regulon dynamic score greater than 0.5 were included. Only the regulons that have more than 5 connections were labeled. The links were colored based on the regulon. **B.** Heatmap indicating the regulon dynamic scores for lineage-regulon combinations for ASD. **C.** Heatmap indicating the-log10(padj) connecting regulons to diseases for Telencephalon: GABA lineage. Only the regulons related to ASD were included. Glut: Glutamatergic neuronal lineage, GABA: GABAergic neuronal lineage, AD: Alzheimer’s disease, AD variants: late-onset (Lo-AD), ADHD: attention deficit hyperactivity disorder, ASD: autism spectrum disorder, BPD: bipolar disease, EPL: epilepsy, MDD: major depressive disorder, SCZ: schizophrenia, SID: Severe intellectual disability, LD: Language disability.

Notably, both cerebellar lineages showed strong associations with a wide range of disorders, underscoring their importance. GABAergic lineages exhibited more disorder associations than glutamatergic lineages, consistent with GABAergic dysfunction as a key mechanism in neurodevelopmental disorders and a potential therapeutic target^64^.

Focusing on ASD (**Figure 5B**), most associations arose from GABAergic lineages, with SOX11, SOX4, and BCL11A strongly linked. The telencephalic GABAergic lineage harbored the largest number of ASD-associated regulons, followed by the cerebellar GABAergic lineage. To further investigate, we examined how these ASD-associated regulons intersected with other disorders within the telencephalic GABAergic lineage (**Figure 5C**). Strikingly, MAFB was associated with all five neurodevelopmental disorders analyzed, while SOX4 and SOX11 showed broader associations spanning both NDDs and additional developmental traits. These findings underscore the pivotal role of neurogenic regulons, particularly within GABAergic lineages, in shaping the molecular landscape of neurodevelopmental disorders and highlight promising avenues for targeted therapeutic exploration.

### Comparative gene regulatory subnetworks of mammalian forebrain neurogenesis

The human telencephalon exhibits specialized and evolutionarily derived molecular, cellular, and anatomical characteristics that distinguish it from other mammalian species^65–67^. However, the complex regulatory architecture governing human telencephalic development is also believed to underlie an increased vulnerability to neurodevelopmental and neuropsychiatric disorders^68,69^. To investigate the evolutionary basis of this regulatory complexity, we analyzed dynamic regulatory subnetworks across the developing telencephalon in human (*Homo sapiens, PCD 35-98*)^19^, rhesus macaque (*Macaca mulatta, PCD 37-78*)^20^, and mouse (*Mus musculus, PCD 7-18*)^21^ aligning timepoints to account for heterochrony (PMID: 23616543) (**Methods**). We identified both telencephalic glutamatergic and GABAergic lineages across all three species (**Figure 6A, Figure S7A**).

**Figure 6.**
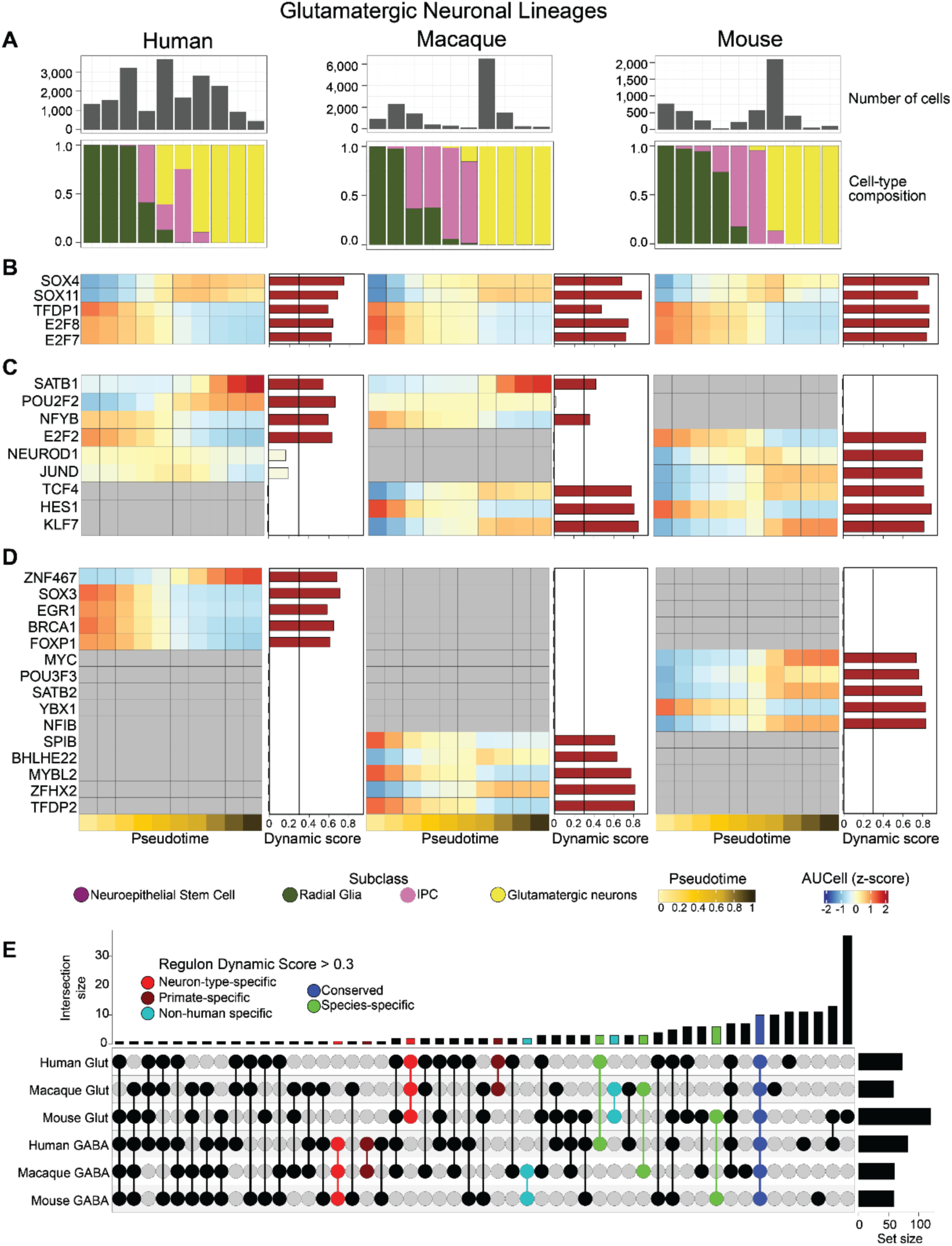
Comparative regulon dynamics across human, macaque, and mouse during telencephalic glutamatergic neurogenesis. **A**. Bar plots showing the number of cells (top) and cell type composition (bottom) across pseudotime bins for human, macaque, and mouse glutamatergic lineages. **B–D.** Pseudotemporal activity profiles of selected regulons grouped by conservation: conserved and highly dynamic across all three species (**B**), pairwise conserved (present in two of three species, **C**), and unique to individual species (**D**). Bar colors denote dynamic scores: red (> 0.3) and tan (< 0.3). **E.** UpSet plot showing overlap among dynamic regulons (score > 0.3). Color codes: blue - conserved across all six glutamatergic lineages, green - species-specific, red - neuron-type-specific, brown - primate-specific, and cyan - non-human-specific.

We first identified the top five conserved dynamic regulons across each species for both glutamatergic and GABAergic lineages (**Figure 6B and Figure S7B**). Notably, many of these conserved regulons are implicated in fundamental biological processes such as proliferation and cell cycle regulation (e.g., E2F family) as well as general neuronal differentiation (e.g., SOX4 and SOX11). These results underscore a core set of genetic programs regulating proliferation and neuronal differentiation that are conserved not only among human brain regions (**Figure S2B**), but also across mammalian species (**Figure 6B and Figure S7B**).

Subsequently, we examined region-enriched dynamic regulon patterns by visualizing the most prominent dynamic regulons through pairwise cross-species comparisons and species-specific visualizations (**Figures 6C, D, and Figures S7C, D**). Interestingly, we identified many species differences in the dynamics of regulons associated with proliferation. The NFYB regulon, essential for neural progenitor maintenance^70^, and MYBL2, a key driver of progenitor population expansion^71^, exhibited pronounced dynamic regulation in primate glutamatergic and GABAergic lineages, respectively (**Figure 6C, Figure S7C)**.

Additionally, we found the BRCA1 and FOXP1 regulons to be highly dynamic in human glutamatergic lineages, with specific enrichment in neural progenitors (**Figure 6D**). The identification of the dynamic BRCA1 regulon in human progenitors underscores derived regulatory networks underlying neural progenitor proliferation, survival, and genomic maintenance (**Figure 6D**). Additionally, the progenitor-enriched FOXP1 regulon represents a candidate for evolutionary specialization in cortical neurogenesis (**Figure 6D**). Expressed in apical and basal RG, FOXP1 regulates pathways essential for progenitor self-renewal and early glutamatergic neuron production^72^. Its human-specific regulon enrichment may support neocortical expansion by maintaining progenitor identity and modulating differentiation timing. Notably, FOXP1 mutations are linked to autism and intellectual disability^73^, suggesting this regulon may underlie early neurodevelopmental disease susceptibility.

To further investigate these comparative GRNs, we analyzed all highly dynamic neurogenic regulons in the telencephalon (dynamic score >0.3) across all species, quantifying the extent of conservation vs. species-specificity (**Figure 6E**). Most regulons were conserved across species and neuronal lineages, underscoring the broad conservation of a core set of neurodevelopmental molecular pathways (**Figure 6E**). Additionally, we identified primate-, non-human-, and neurotransmitter-phenotype-specific subsets (**Figure 6E, Figure S7E)**. Together, this comparative regulon analysis provides a mechanistic framework for elucidating how both conservation and divergence of dynamic developmental gene regulatory programs contribute to specializations in telencephalic development, function, and disease susceptibility.

### Developmental regulatory network resource of mammalian neurogenesis

So far, we have investigated neurogenic lineages across multiple mammalian species and human brain regional hierarchies, highlighting both shared and region-specific features. To unify our lineage analyses, we performed hierarchical clustering on the dynamic regulon scores of all 18 human neurogenic lineages (**Figure 7A**). The resulting lineage dendrograms revealed biologically coherent patterns: all three midbrain lineages clustered together, reflecting their cohesive regional identity; pons and medulla lineages grouped within the hindbrain branch, whereas cerebellar lineages formed a distinct cluster, indicating regulatory divergence within the hindbrain. In the forebrain, telencephalic glutamatergic and diencephalic lineages clustered closely with the broad forebrain lineage, whereas telencephalic GABAergic lineages formed a separate cluster, suggesting that neuronal lineage identity exerts a stronger influence on the dynamic regulatory programs than regional similarity in the developing telencephalon.

**Figure 7:**
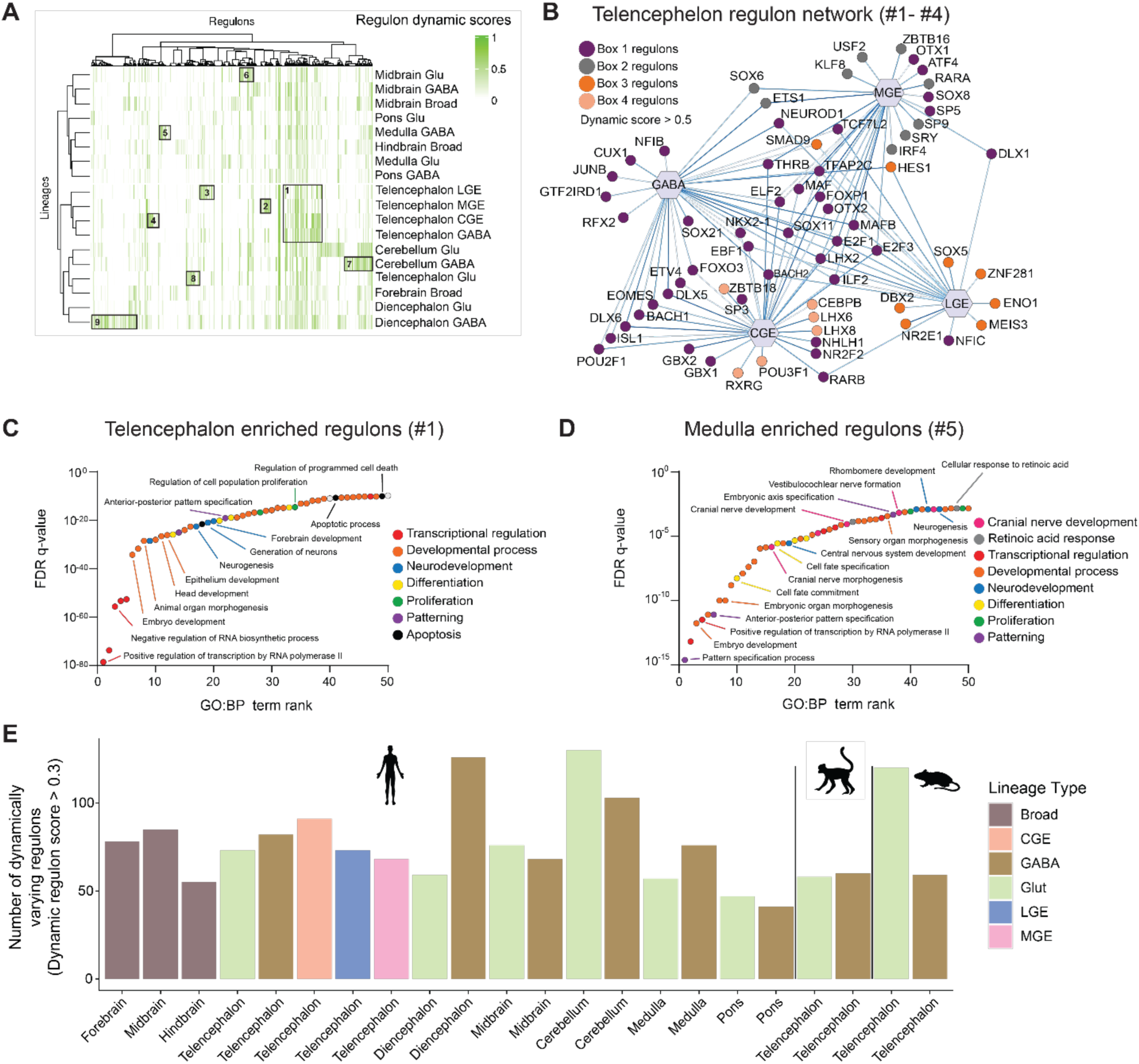
Developmental regulatory network resource of mammalian neurogenesis. **A.** Heatmap indicating the dynamic regulon scores for the 18 human neurogenesis lineages. The dendrograms are obtained using hierarchical clustering. Boxes are drawn around groups of regulons enriched in specific regions (box #1, telencephalon GABA lineages) or specific lineages (box #2, MGE; #3, LGE; #4, CGE; #5, medulla GABAergic; #6, midbrain glutamatergic; #7 cerebellum GABAergic; #8, telencephalon glutamatergic; #9, diencephalon GABAergic). **B.** A bipartite network connecting lineages and regulons highlighted in boxes 1-4 in **Figure 7A**. Nodes are colored by their association with the boxes highlighted in **Figure 7A**. Box 1 - regulon cluster conserved in all 4 telencephalic GABAergic lineages. Box 2-4: regulon clusters that show specificity to MGE, LGE, and CGE lineages, respectively. **C-D**. Waterfall plots depicting Gene ontology (GO: Biological processes) terms enriched in Box 1 (**C**), and Box 4 (**D**). **E.** A barplot showing the number of dynamically varying regulons (i.e., dynamic score > 0.3) in all 22 neurogenesis lineages. Lineages are grouped according to their species and colored according to their regional hierarchy.

The simultaneous clustering of regulons revealed striking regulatory signatures. For instance, telencephalic GABAergic-enriched regulon groups (boxes 1-4) represent conserved (box 1) and developmental origin-specific (boxes 2-4) regulons (**Figure 7B**). Gene set enrichment analysis of box 1 regulons revealed key neurodevelopmental pathways, including transcription regulation, anterior-posterior patterning, proliferation, differentiation, apoptosis, and forebrain development (**Figure 7C**). These findings align with established developmental programs fundamental to telencephalic GABAergic neurons, encompassing precursor proliferation, differentiation into immature interneurons, and intrinsic, developmentally programmed apoptotic pathways.

Furthermore, we identified a distinct cluster of regulons highly enriched in the medulla GABAergic neuronal lineage (box #5) (**Figure 7A**). Similarly, these medulla-enriched dynamic regulons were significantly associated with neurodevelopment-related biological processes, including transcriptional control, cellular proliferation, and the specification of developmental axes (**Figure 7D**). Unique to this cluster, however, was a pronounced enrichment for processes specific to medulla development, such as cranial nerve formation, morphogenesis, and rhombomere segmentation. Among these, ‘pattern specification process’ emerged as the most significantly enriched GO: BP term, underscoring the critical role of dynamic regulons in the precise patterning and segmentation of the developing medulla. Collectively, these results underscore the advantage of a dynamic regulon-based lineage analysis framework in uncovering key neurodevelopmental regulatory nodes.

This approach enabled the identification of principal transcriptional regulators governing both general neurogenic processes—such as proliferation and differentiation—and region-specific developmental programs, including forebrain versus hindbrain patterning. Our findings demonstrate that regulon-directed analyses not only facilitate robust feature selection but can also provide mechanistic insights into the critical biological drivers driving complex neurodevelopmental trajectories. This framework represents an invaluable tool for dissecting the transcriptional architecture that underlies the spatial and temporal regulation of neurogenesis. As a summary of our findings, we tabulated the number of dynamic regulons (dynamic scores > 0.3) for all 22 lineages analyzed in this study (**Figure 7E**).

## DISCUSSION

This study delivers a dynamic and systems-level analysis of gene regulation during early mammalian neurogenesis, revealing how temporal, spatial, and cell-type-specific transcriptional programs shape brain development. Our computational analysis discovers temporally resolved, biologically interpretable regulatory subnetworks, capturing the regulatory logic of developmental trajectories with unprecedented precision. Applying this approach, we mapped 18 human neurogenic lineages across three hierarchical levels spanning the whole brain and four non-human lineages, creating the most comprehensive regional and cross-species resource of developmental GRNs to date. This resource reveals how transcription factors and their targets, organized into regulons and gene modules, coordinate neurogenesis across diverse brain regions and species, advancing both the conceptual and methodological foundations for studying dynamic gene regulation.

By integrating trajectory inference, pseudotime analysis, and regulatory network modeling, we uncover both conserved and context-specific transcriptional regulators that define broad neurogenic processes as well as molecular heterogeneity within discrete brain regions and cell types. Our findings validate established regulators as core components of neuronal differentiation, while also uncovering previously uncharacterized and highly dynamic regulons, including candidates with restricted regional or lineage enrichment such as TEAD2 and ZIC3. These findings illustrate the complex, multilayered regulatory architecture underlying neurotransmitter identity, subregional specification, and dynamic transitions along neurogenic trajectories.

Cross-species comparisons further uncovered the evolutionary conservation of core proliferation and differentiation programs along with the emergence of lineage-or species-enriched regulons that may contribute to derived features of primate brain development. Together, our findings reveal an extraordinary degree of precision and complexity in the regulatory mechanisms that drive nervous system formation from its earliest embryonic stages, highlighting the dynamic interplay between genetic programs and developmental cues that shape neuronal identity and connectivity.

Furthermore, this resource provides novel insights into the molecular etiology of neurodevelopmental disorders by mapping the dynamic regulatory landscapes associated with disease-linked gene modules and identifying cell lineages with heightened genetic risk. We correlated functional gene modules to neurodevelopmental disorders via GWAS traits and found links between individual GABAergic neurogenic lineages, dynamic gene modules, and neurodevelopmental disease risk. Additionally, using known disease transcriptomic signatures and calculating their dynamic association to individual neurogenic lineages, we found telencephalic enrichment of neurodevelopmental disorder disease gene signatures, providing guidelines for future exploration of disease etiology.

To facilitate further exploration of this resource by the scientific community, all inferred GRNs, dynamic scores, and related supplementary data tables are provided. We have developed an interactive online platform (https://daifengwanglab.shinyapps.io/devGRNDB/) (**Figure S8**) that enables direct comparison of lineages and regulatory networks, enhancing accessibility and utility for future research. Additionally, the complete computational framework is made publicly available via GitHub for reproducibility and extension of our findings.

Our results further substantiate the advantage of regulon-based analyses over single-gene approaches, offering enhanced sensitivity and mechanistic insight into transcriptional regulatory dynamics from sparse single-cell data. Collectively, this work not only expands our understanding of lineage-and region-specific neurodevelopmental programs but also lays a foundation for future investigations linking dynamic regulatory network perturbations to specific cellular, anatomical, and functional outcomes in both health and disease. The presented GRN resource and accompanying online platform serve as valuable resources for the neuroscience and genomics communities, providing a robust framework for interrogating the molecular basis of brain development, evolution, and disease.

## ACKNOWLEDGEMENTS

This work was supported by the National Institutes of Health (RF1MH128695, R01AG067025 to D.W.; 1R01HD106197, UM1MH130991 to A.M.M.S.; and P50HD105353 to Waisman Center). Further support was provided by the National Science Foundation Career Award (2144475 to D.W.), SFARI pilot grant 971316 (X.Z., A.M.M.S. and D.W.), National Institutes of Health 1F30MH140382-01, the Medical Scientist Training Program T32 (GM140935), and the Morse Society Fellowship to R.D.R.

## AUTHOR CONTRIBUTIONS

Conception: DW Conceptualization: KHA, DW

Methodology: KHA, RDR, JS, PK, DW

Formal analysis: KHA, RDR, JS, PK, JC, SA, CG, AMMS, DW

Writing: KHA, RDR, JS, PK, JC, SA, XZ, AMMS, DW

Visualization: KHA, RDR, JS, PK, JC, SA

Supervision: XZ, AMMS, DW

Funding acquisition: DW, AMMS

All authors read and approved the final draft of the paper.

## RESOURCE AVAILABILITY

The datasets used in the current analysis are publicly available at Braun et al.^19^ (Human), La Manno et al.^21^ (Mouse), and Micali et al.^20^ (Macaque). The neurogenic gene regulatory resource can be interactively explored at https://daifengwanglab.shinyapps.io/devGRNDB/ (**Figure S8**). The computational codes for the proposed framework can be accessed at https://github.com/daifengwanglab/devGRNDB/. All other data are available in the main article or supplemental information. Further information and requests for resources should be directed to the corresponding authors.

## SUPPLEMENTAL INFORMATION

Supplementary Figures 1-8.

Supplementary Tables 1-7.

## METHODS AND MATERIALS

### Computational framework for dynamic regulatory networks in development

We developed a computational framework to identify gene regulatory subnetworks associated with any given biological process. This framework integrates existing tools for lineage tracing, pseudotime estimation, and gene regulatory network (GRN) inference. Importantly, while we provide recommended tools within this manuscript, users can flexibly incorporate alternative bioinformatic methods that perform equivalent tasks and are best suited to their dataset.

Our framework comprises four key steps:

Step 1: Input data preparation

The input to the framework is single-cell or single-nucleus gene expression data annotated with cell type, relevant to the biological process under investigation. Although we showcased our framework in studying developmental trajectories, it is equally applicable to disease progression analyses. Therefore, metadata must include either temporal information or clinical indicators of progression to enable pseudotime inference.

Step 2: Lineage inference and pseudotime estimation

In this step, cells are ordered along a biological continuum (e.g., differentiation or disease progression). When multiple lineages are present, such as distinct cell fates in neurogenesis, we recommend using cell fate prediction algorithms such as CellRank, which calculate fate probabilities delineating lineage-specific cell subsets.

Step 3: Construction of lineage-specific GRNs

Next, GRN inference is conducted separately for each lineage, based on the subset of cells assigned to that lineage. These subsets may contain multiple cell types and cellular states, allowing for inference of consensus GRNs. While modeling the dynamic activity of each transcription factor (TF)–target gene (TG) pair over pseudotime is informative, it is often computationally intensive and highly sensitive to the noise inherent in single-cell expressions. Therefore, we focus on biologically meaningful regulatory subnetworks such as gene modules or regulons (i.e., TFs and their target genes), which improve both computational tractability and robustness.

Step 4: Quantification and prioritization of dynamic regulatory subnetworks for lineages.

Finally, we assess the activity of each regulatory subnetwork across individual cells using an enrichment-based scoring approach. In this study, we apply an enrichment-based method (i.e., AUCell^11^, UCell^74^), which ranks genes by expression within each cell and computes an enrichment score for the subnetwork gene set. These scores can then be utilized to explore their association with the biological trajectory. As subnetwork dynamics may exhibit complex patterns such as monotonically increasing or decreasing, biphasic, or static, a robust statistical approach is required to quantify their association. To this end, we use Moran’s I, a spatial autocorrelation metric previously employed in Monocle3 to detect trajectory-associated genes. While Monocle3^75^ utilizes graph-based spatial coordinates, our approach defines a multi-dimensional pseudo-space derived from pseudotime. To account for inherent uncertainty in pseudotime inference, we implement an ensemble approach: multiple pseudotime estimations are made by varying root cells and randomly subsampling gene features. These pseudotime values are then integrated to form a multi-dimensional pseudo-space, which provides a more robust foundation for computing spatial autocorrelation.

A key strength of this framework is its adaptability to diverse and emerging computational tools. Given the continuously growing number of methods for trajectory inference^75–77^, and GRN reconstruction^11,78–81^, our framework allows users to flexibly integrate current or future tools at each step.

Our computational framework can be readily extended to incorporate additional modalities beyond gene expression. For instance, joint profiling of gene expression and chromatin accessibility enables the construction of more comprehensive gene regulatory networks. In such cases, integrative computational tools such as Pando^82^ or SCENIC+^10^ can be employed. Notably, SCENIC+ includes built-in functionality to compute AUCell scores based on both gene expression and chromatin accessibility for inferred regulons. These outputs can be seamlessly integrated into our framework to evaluate regulon activity and its association with the biological process of interest.

### Construction of a gene regulatory network atlas for neurogenesis Input data preparation

#### scRNA-seq preprocessing

We used single cell data from Braun et al.^19^, which had studied the developing human brain during the first trimester. ∼1.67 million single cell RNA sequencing data from 26 brain specimens, ages spanning from 5 to 14 postconceptional weeks (pcw), were taken for the study. The dataset dropped down to ∼1.2 million cells when isolated only to the neurogenesis-related cell types: Neurons, Neuroblasts, Neuronal IPC, and Radial glia. Based on the brain region information present in the original dataset, it was divided into three broad regions: forebrain, midbrain, and hindbrain, and six subregions: telencephalon, diencephalon, midbrain, pons, medulla, and cerebellum. Each brain region dataset was processed independently to obtain region-specific cell subtype annotations. We used SCANPY^83^ (v1.11.0) for data preprocessing, clustering, and cell annotation.

#### Celltype annotation

Cell type annotations were performed independently for each of the six brain subregions to enhance annotation accuracy. The dataset contains cells from different brain regions collected from donors at different ages. Hence, for a given brain region, both sample age and the cellular identity (i.e., cell type) contribute to the patterning of cellular transcriptomic profiles. Given the pronounced transcriptomic shifts in early fetal cells, calculating highly variable genes (HVGs) across the entire dataset could obscure biologically relevant signals from early developmental stages^20,84^. To preserve age-specific transcriptomic features, the top HVGs were identified independently for each sample (i.e., age group). The union of these HVGs was then used for downstream analysis. Then we performed normalization, log-transformation, and scaling. The top 50 principal components were calculated using the HVGs, and the samples were integrated using the harmony_integrate method^85^ available in Scanpy external API (v1.11.0). Then nearest neighbors were calculated using a local neighborhood of 20 with harmony components (X_pca_harmony). Next, Leiden^86^ clusters were obtained with a resolution of 2.0 and manually annotated using the marker genes provided in **Table S2**.

#### Pseudobulk calculations

Given the computational expense of processing ∼1.2 million cells and the inherent noise in single-cell data, we aggregated cells into pseudo-bulk profiles. The pseudobulk cells were obtained by aggregating the gene expression of 10 transcriptomically similar cells (i.e., 10 closest neighboring cells). To account for variations in age, brain region, and cell type across the dataset, pseudobulk calculations were performed iteratively within subsets of cells sharing the same region, age, and cell type. This ensured accurate metadata assignment for each pseudobulk cell and preserved biologically meaningful differences across developmental and spatial contexts. Given the substantial shifts in cell type composition with age and region, we maintained the relative proportions of each cell type within the pseudobulk sampling to minimize artificial biases. The pseudobulk generation followed these steps:

1. Isolation of cells from a defined age group, brain subregion, and cell type
2. Identification of 10 nearest neighbors using a k-nearest neighbors algorithm based on the top 50 principal components
3. Estimation of pseudobulk count using a 10:1 ratio (i.e., 10 single cells per pseudobulk)
4. Random selection of root cells to seed pseudobulk profiles
5. Aggregation of gene expression from each root cell’s nearest neighbors to construct final pseudobulk profile

Due to the limited availability of cells in pons, the number of pseudobulk cells was determined by the ratio of 5:1.

These pseudobulk cells were used for downstream analysis, including the steps of our proposed computational framework. This procedure was implemented on two levels of regional hierarchies (**Figure 1B**). In the broad regions (i.e., forebrain, midbrain, hindbrain), pseudobulk cells were calculated using the cell class annotations from the original publication [PMID: 37824650]. For subregions (i.e., telencephalon, diencephalon, etc.), our cell subclass annotations were used. The same subregion pseudobulk cells were used in the GABAergic subtype neuronal lineages (i.e., CGE, LGE, etc.). Details on the pseudobulk samples are provided in **Table S2**.

#### Processing non-human samples

To facilitate precise cross-species comparison of neurodevelopment, we quantitatively aligned developmental stages across all three species, correcting for telencephalic heterochrony^87^. Subsequently, we isolated cells from developing mouse^21^ and macaque^20^ telencephalon datasets at ages corresponding to the aligned human developmental timepoints (mouse: 7–18 post-conception days [PCD]; macaque: 37–78 PCD). Mouse cells were annotated using the same procedures detailed in the Celltype annotation section (see **Table S2** for marker genes), whereas original cell annotations provided with the macaque dataset were retained. For both mouse and macaque, pseudobulk calculations were performed as described for the human dataset.

### Lineage and pseudotime inference

#### Pseudotime inference

We performed pseudotime inference using pseudobulk data to capture lineage trajectories, while reducing noise from single-cell variability and sparsity. First, we selected the top 5000 HVGs across pseudocells using SCANPY^83^ (v1.10.4) (sc.pp.highly_variable_genes). HVG selection reduced biological and technical noise and improved computational efficiency for downstream analyses. Next, we scaled the data (sccanpy.pp.scale) and performed dimensionality reduction using principal component analysis (PCA) (scanpy.tl.pca). Following PCA, we corrected batch effects between pseudocells using Harmony^85^ integration (scanpy.external.pp.harmony_integrate). We subsequently constructed a neighborhood graph using the Harmony-integrated PCA space. Next, we ran Palantir^76^ (v1.3.6) on the Harmony-integrated PCA space, using the previously computed kNN graph as an adjacency constraint (use_adjacency_matrix=True). The pseudotime was inferred with the selected root and terminal cells passed to scanpy.external.palantir_results (specific subtypes and pseudocell IDs are listed in **Table S2**), and the resulting pseudotime values were subsequently used as input for CellRank to assign lineage memberships and compute fate probabilities.

#### Lineage membership assignment

To infer directed developmental trajectories and lineage fate probabilities, we applied Cellrank^77^ (v2.0.6) with a pseudotime-based transition model. We constructed a PseudotimeKernel (cellrank.kernels.PseudotimeKernel) using the pseudotime values computed from Palantir^76^. The kernel modifies the neighborhood graph such that cell transitions are biased towards forward propagation along increasing pseudotime, thereby guiding directionality into the Markov chain without requiring RNA velocity data. We used the default hard thresholding scheme. We then used the GPCCA estimator (cellrank.estimators.GPCCA) to decompose the transition matrix into macrostates. To refine lineage-specific inference, we manually designated terminal macrostates corresponding to mature neuron subclasses (**Table S2**). We then computed fate probabilities to the curated terminal macerostates using GPCCA.compute_fate_probabilities. Each pseudocell was assigned to the terminal lineage for which it had the highest fate probability.

#### Lineage-specific subsetting and pseudotime refinement

After assigning pseudocells to terminal lineages, we partitioned the dataset into lineage-specific subsets by selecting cells with the highest fate probabilities for each respective lineage. This partitioning strategy allows us to isolate lineage-committed cells and focus our analysis on lineage-specific differentiation trajectories. To improve the robustness of pseudotime inference within each lineage, we performed iterative Palantir pseudotime analysis with multiple random initializations. For each iteration, we recomputed the k-nearest neighbor (kNN) graph using the Harmony-integrated PCA embedding with different random seeds and selected a distinct root pseudocell from the same progenitor cell population. We conducted four independent iterations per lineage, with each iteration using a unique root cell selected from within the progenitor subset, ensuring that our pseudotime estimates were not dependent on the choice of a single starting cell.

### Quantification and prioritization of dynamic regulatory subnetworks for lineages

#### Gene regulatory network inference for neurogenic lineages

In the current study, we have utilized the SCENIC^11^ framework implemented in pySCENIC^79^ (v0.12.1+8.gd2309fe) to infer the gene regulatory networks. First, the log-normalized gene expression data were used to infer a co-expression network by connecting transcription factors (TFs) to potential target genes (TGs) based on gene-gene correlations using GRNboost2^81^. The species-specific TF lists provided in the pySCENIC package [https://resources.aertslab.org/cistarget/tf_lists/] were utilized. Then these target genes were further refined by examining the transcription factor binding motif enrichment in the upstream of the genes using RcisTarget. Due to the unavailability of macaque-specific databases (i.e., TF list, cisTarget database), human databases (‘hg38’) were incorporated for macaque GRN inference steps. Mouse GRNs were inferred using Mus musculus - mm10 datasets provided in the pySCENIC package (https://resources.aertslab.org/cistarget/databases/, https://resources.aertslab.org/cistarget/motif2tf/). This provides a directed gene regulatory network connecting transcription factors to their target genes. All 22 GRNs inferred for the 22 neurogenic lineages are provided in **Table S3**.

#### Consensus feature selection for GRN inference

In the current study, we have explored GRNs for 22 lineages covering three species and multiple brain regions at different hierarchical levels. Since we provide a regulatory network atlas that allows cross-comparison, we utilized a consensus gene set that represents all 22 lineages. This was obtained by independently estimating the top 5000 highly variable genes for each lineage. This led to 11699 unique highly variable genes.

#### Definition of regulatory subnetworks

In this analysis, we examined two types of regulatory subnetworks: regulons and co-regulatory gene modules (**Table S1**), to evaluate their relevance to neurogenesis based on dynamic scores. A regulon is defined as a set consisting of a transcription factor (TF) and all of its predicted target genes (TGs). To ensure statistical robustness, only the TFs with at least five associated TGs were considered in our analysis.

The second type of regulatory subnetwork is co-regulatory gene modules, which capture patterns of shared regulation across genes. Since our lineage-specific gene regulatory networks (GRNs) define only TF-to-TG relationships, they do not capture interactions among non-TF genes. Therefore, these GRNs cannot be used directly to infer gene co-regulation modules. To address this, we constructed a co-regulatory network by estimating the Jaccard index between the sets of regulators (TFs) associated with each gene pair.

For two genes 𝑔_i_ and 𝑔_j_, let 𝑅(𝑔_i_) and 𝑅(𝑔_j_) denote their respective sets of regulating TFs. The Jaccard index is defined as: 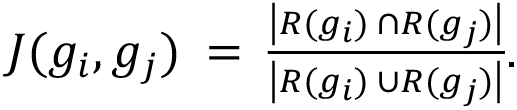 This index ranges from 0 to 1, where 𝐽 = 0 indicates no shared regulators and 𝑗 = 1 indicates an identical set of regulators. The resulting Jaccard similarity scores form an adjacency matrix, 𝐴, representing the co-regulatory network. To identify gene modules, we first computed a dissimilarity matrix as 1 − 𝐴. This matrix was then subjected to hierarchical clustering using the “average” linkage method via the hclust function in R (v4.5.0). Gene modules were identified by cutting the resulting dendrogram using the cuttreeDynamic function from the WGCNA^88^ package (v1.73), with a minimum module size threshold set to 100 genes.

### Prioritization of regulatory subnetworks

#### Estimation of subnetwork activity

Once the regulatory subnetworks were identified, per-cell regulatory activity scores were inferred using the gene sets in each subnetwork (i.e., in gene modules, all the genes in the gene module). In the current study, we utilized the AUCell^11^ approach (v1.30.1), which calculates how highly the genes in a given gene set are expressed compared to a ranked list of all the genes expressed in a cell via the Area Under the Curve (AUC) method. We utilize this score as an indicator of how active a regulatory subnetwork is in a given cell. Alternatively, any gene signature scoring method, such as UCell, could be used instead of AUCell. To reduce the influence of noise, we excluded regulons that exhibited non-zero regulon activity in less than 1% of cells within the corresponding lineage.

#### Dynamic score calculation

We utilized Moran’s I statistic to investigate the global level association of the regulatory activity with the inferred pseudotime. We used the Moran.I function in ape package (v5.8-1) with the arguments, scaled = TRUE and alternative = ‘greater’. To account for uncertainties in pseudotime estimation, we computed five versions of pseudotime for the same lineage (see Step 2: Lineage and pseudotime inference). These pseudotimes were used to construct a five-dimensional pseudo-space representing relative distances among cells based on pseudotime. From this, we derived a Euclidean distance matrix, whose inverse was used as the weight matrix input for the Moran.I function. Adjusted p-values were calculated using the p.adjust function in R, with ‘BH’-Benjamini & Hochberg method^89^ applied for multiple testing correction. We have utilized the dynamic score cutoff of 0.3 to identify dynamically varying regulons, except for **Figure S2E** and the disease overrepresentation analysis (**Figure 6**), where cutoff values of 0.4 and 0.5 were used, respectively. This was primarily to improve the readability of the visualizations. Detailed lists for the corresponding analyses are provided in **Tables S4,5, and 7**.

#### TF gene expression vs. regulon activity

Here, we compared the possibility of using TF gene expression itself for the dynamic score instead of using regulons. The Moran’s I statistic has been previously utilized in Monocle3, a well-known trajectory inference technique, to investigate differentially expressed genes along the trajectory—that is, the dynamic association between gene expression and trajectory. Thus, as an alternative hypothesis, we calculated dynamic scores using TF gene expressions and compared them with regulon dynamic scores (**Figures S1A–C**). We followed the same procedure described in “Dynamic Score Calculation” to estimate Moran’s I statistic. We compared the two dynamic scores using a paired Wilcoxon signed-rank test.

### Functional Analysis

#### Sensitivity analysis for GRN inference methods

We performed a sensitivity analysis for our computational framework by using three GRN inference approaches, namely, GRNBoost^81^, SCENC^79^ (v0.12.1+8.gd2309fe), and SCODE^80^. The GRNBoost and SCENIC inferences were done as explained in the Gene regulatory networks and subnetwork inference section. Only the top 10% TF-TG links by edge weight for SCODE and GRNBoost were used for this analysis. SCODE incorporates pseudotime as an input for the ordinary differential equation based algorithm to infer gene regulatory networks (GRNs). We tested the three GRN approaches using single-cell gene expression data from the caudal ganglionic eminence (CGE) of the developing telencephalon. To make it comparable, we built GRNs for the top 5000 HVGs and kept the same set of TFs across methods.

In the SCODE, we optimized the model by evaluating different values for the latent dimensionality parameter 𝐷 and selected the value that minimized the reconstruction error of the expression data. Then, using the optimized D value, we inferred a GRN using SCODE.

#### Capturing putative regulatory interactions

The three GRNs obtained were then input into our framework. We first identified the regulons in each GRN by isolating the TFs that have more than 5 TGs. Then we estimated their regulon activities using AUCell. Dynamic scores were then calculated using the pseudotime values obtained for the Telencephalon CGE lineage. As expected, the three GRNs provided different TF-TG links and resulted in different lists of regulons (SCENIC - 50, GRNBoost - 301, SCODE - 179). Out of these regulons, 12 regulons were conserved in all three methods, and 12 and 50 regulons were overlapped between SCODE-SCENIC and GRNBoost and SCENIC, respectively (**Figure S3A**). The pairwise comparisons were made against SCENIC results (**Figures S3B-C**). We then estimated pairwise F-1 scores to quantify the sensitivity of the GRN inference method in our computational framework in predicting dynamically varying regulons (i.e., dynamic score > 0.3). With that, the SCODE-SCENIC and GRNBoost-SCENIC comparisons provided F-1 scores of 0.94 and 0.96, respectively. The three GRNs, their regulon dynamic scores are provided in **Table S2**.

#### Disease enrichment and gene module enrichments

Gene modules for CGE, LGE, and MGE that were obtained using the method described in Definition of regulatory subnetworks were used to do disease enrichment. We then performed linkage disequilibrium (LD)-aware enrichment analysis of GWAS traits using MAGMA (Multi-marker Analysis of GenoMic Annotation)^57^. Summary statistics from multiple GWAS were curated, and SNPs with nominal p-values (p ≤ 0.05) were annotated to genes based on human genome build GRCh38 coordinates from NCBI. Enrichment analyses utilized reference LD structure from European ancestry samples in Phase 3 of the 1000 Genomes Project. The sample size parameter N was set according to values reported in the respective GWAS studies: schizophrenia^90^ (N = 76,755), autism spectrum disorder^91^ (N = 46,350), bipolar disorder^92^ (N = 41,917), and epilepsy^93^ (N = 29,944).

### Lineage and neurological disorder associations

#### Curation of disease risk genes for neurological disorders, cortical malformations, and developmental traits

We have conducted an exploratory investigation to find dynamic associations between known neurological disorders with the 18 human neurogenesis lineages. We first curated the genes associated with selected neurological disorders, cortical malformations, and developmental traits. The disease risk genes were curated from multiple resources (DisGeNet (https://disgenet.com/, accessed [Feb, 2023], SFARI (https://gene.sfari.org/, accessed [July, 2025]), and SZDB database (http://szdb.org/ accessed [July, 2025] and Mato-Blanco et al.^94^). The neurological disorders include epilepsy (EPL), autism spectrum disorder (ASD), attention deficit hyperactivity disorder (ADHD), neuroticism (NEUROT), major depressive disorder (MDD), Alzheimer’s disease (AD), Familial-Alzheimer’s disease (FAD), Late Onset Alzheimer’s disease (LoAD), Early Onset Alzheimer’s disease (EoAD), Focal Onset Alzheimer’s disease (FoAD), bipolar disease (BPD), schizophrenia (SCZ), Early Onset schizophrenia (EoSCZ), anorexia nervosa (AN), Parkinson’s disease (PD) and young onset Parkinson’s disease (YoPD). Among the developmental traits, we considered severe intellectual disability (SID), dyscalculia, dysgraphia, performance anxiety, central auditory processing disorder (cAPD), non-verbal learning disorder, language disorders (LD), intelligence, and developmental coordination disorder (DCD). Cortical malformations include macrocephaly, macrocephaly at birth, microcephaly, cobblestone, heterotopia, hydrocephalus, focal cortical dysplasia and the mammalian target of rapamycin (FCDandmTOR), polymicrogyria, and rare malformation of cortical development (rareMCD). From the SFARI gene list, only the genes with a gene score of 1 or 2 were selected. For Schizophrenia, gene lists from the SZDB database were filtered based on the Pascal scores 1 to 6. All the disease risk genes and their sources were tabulated in **Table S6**.

#### Overrepresentation of regulon in disease genes

With the intention of identifying neurogenic regulators (i.e., dynamically varying regulons) that could potentially be disrupted due to the aforementioned neurological disorders, we conducted overrepresentation analysis of inferred regulons in disease genes using a hypergeometric test. The hypergeometric test p-values were then adjusted for multiple testing using Benjamini-Hochberg method and retained only the statistically significant associations (i.e., adjusted p-value < 0.05, **Table S6**). This allowed us to link regulons with disease risk genes and ultimately to specific neurogenic lineages via dynamic regulon scores. **Figure 6A** depicts these lineage-regulon-disease associations using a sankey plot (with a dynamic regulon score cutoff of 0.5).

#### Hierarchical Clustering of Lineage-Regulons

We performed hierarchical clustering on regulon dynamic scores for regulons across lineages to identify lineage-specific regulon clusters. Here, we first computed pairwise dissimilarities between lineages and regulons using “correlation” based distance. To do this, we used the hclust() and proxy() (v0.4-27) functions in R. Then, we applied agglomerative clustering with Ward’s D2 linkage independently to both axes, yielding dendrograms that were cut into different regulon clusters (10, 40) to capture finer regulon clusters. To do this, we used the Heatmap() function from the ComplexHeatmap (v2.24.1) package in R.

#### Regulon selection for pseudotemporal regulon activity heatmap visualization

We presented three categories of regulon sets for visualizing pseudotemporal regulon activity: (1) those conserved across all lineages, (2) those conserved between two lineages (i.e., pairwise conservation), and (3) those unique to a specific lineage. These classifications and their rankings were primarily based on regulon dynamic scores. For the conserved category, we first identified regulons present across all lineages and then ranked them according to their maximum dynamic score observed among the lineages. In the pairwise conserved category, regulons shared between any two lineages were selected, and the top-ranked regulons for each lineage pair were determined. Finally, in the lineage-specific category, we retained regulons exclusive to a single lineage and ranked them based on their dynamic scores.

#### Gene ontology over-representation analysis

We conducted GO over-representation analyses at two different contexts. The over-representation analysis related to all gene co-regulatory modules was performed using the enrichGO() function available in clusterProfiler^95^ (v4.16.0). Multiple hypothesis testing was performed using the Benjamini-Hochberg correction method and filtered to include the Gene Ontology: Biological Processes, which had an adjusted p-value less than 0.01. Human annotations were incorporated using org.Hs.eg.db (v3.21.0). All the genes included in the corresponding co-regulatory network were provided as the background gene set.

Gene set enrichment analysis (GSEA) for hindbrain conserved regulons (**Figure S5F**) and region-enriched regulons (**Figures 7C-D**) was performed using the Molecular Signatures Database (MSigDB; msigdbr v7.5.1) C5 ontology gene set (Homo sapiens; category: Gene Ontology Biological Processes [GO: BP])^96,97^. Statistical significance was assessed by controlling for multiple hypothesis testing, with the false discovery rate (FDR) q-value reported.

#### PHATE and UMAP visualizations

To visualize cell classes within brain sub-regions, we performed dimensionality reduction using both UMAP and PHATE. For UMAP^98^ (Uniform Manifold Approximation and Projection), we used the implementation in the Scanpy toolkit (scanpy.pl.umap version 0.5.7) to project cells into a 2D space based on neighborhood graph relationships derived from the processed gene expression data. Cells were colored by their annotated cell classes to highlight the lineages. In parallel, we applied PHATE^99^ (Potential of Heat-diffusion for Affinity-based Transition Embedding), a non-linear manifold learning method designed to capture continuous developmental trajectories while preserving global data structure. We applied PHATE (v1.0.11) using phate.PHATE(knn=5, decay=20, t=150) on Harmony-corrected PCA embeddings.

