## Supplemental Information for "Dynamic landscapes of gene regulatory networks in early mammalian neurogenesis: Insights into brain evolution and disorder risk"

### SUPPLEMENTARY TABLES

**Table S1:** Glossary of terms

| Term | | Description |
|---|---|---|
| Subnetworks | | A subset of a gene regulatory network, defined by a specific network topology or similarity criterion. |
| Subnetworks | Regulons | A subnetwork representing the transcription factor (i.e., regulator) and its direct target genes. The regulons are named after the corresponding transcription factor. |
| | Co-regulatory modules | Gene cluster linking genes coregulated by the same TFs, weighted by the similarity of their regulators (**Methods**) |
| Single cell level subnetwork activity | | Per-cell enrichment (AUCell)[1] values that quantify the activity of a gene set of a given regulatory subnetwork |
| Lineage level dynamic score of the subnetwork | | A measure of subnetwork activity dynamics, defined as Moran's I (spatial autocorrelation) of subnetwork activities of the cells from the trajectory. |
| Dynamically varying subnetworks/ Dynamic subnetworks | | Regulatory subnetworks with dynamic scores above a cutoff (**Methods**) |

**Table S2:** Data preprocessing, cell type annotation, lineage inference, and sensitivity analysis

**Table S3:** Gene regulatory networks obtained for the 18 human, 2 macaque and 2 mouse neurogenic lineages

**Table S4:** Regulon dynamic scores inferred for the 18 human, 2 macaque and 2 mouse neurogenic lineages

**Table S5:** Co-regulation gene modules, membership, module dynamic scores and GO: BP enrichment terms for all 18 human neurogenic lineages

**Table S6:** Regulon-disorder associations for the human subregion ($2^{nd}$ hierarchy) neurogenic lineages

**Table S7:** Extended summary tables for Figures 2,3,4, and 6.

### SUPPLEMENTARY FIGURES

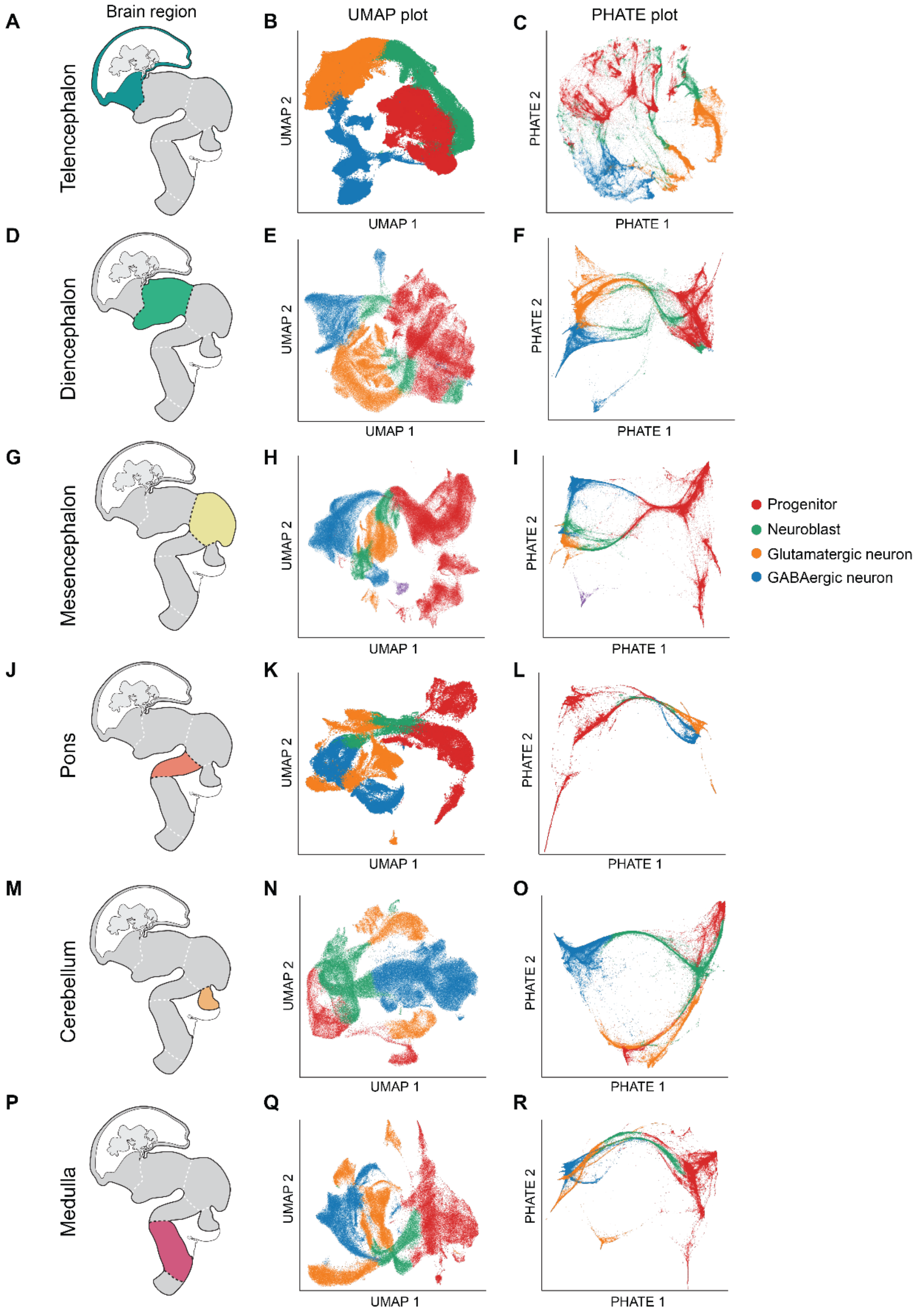

**Figure S1. Human brain regions and cell type visualizations.** Schematic visualization of select human brain regions analyzed in this study (left column), UMAP plot of cell types in corresponding brain regions (middle column), and PHATE[2] plot (right column of cell types in corresponding brain regions for the telencephalon (**A-C**), diencephalon **(DF**), mesencephalon (**G-I)**, pons (**J-L**), cerebellum, (**M-O**), and medulla (**P-R**). Cell types are colored by cell type: neural progenitor (red), neuroblast (green), glutamatergic (orange), GABAergic neuron (blue), serotonergic neuron (purple).

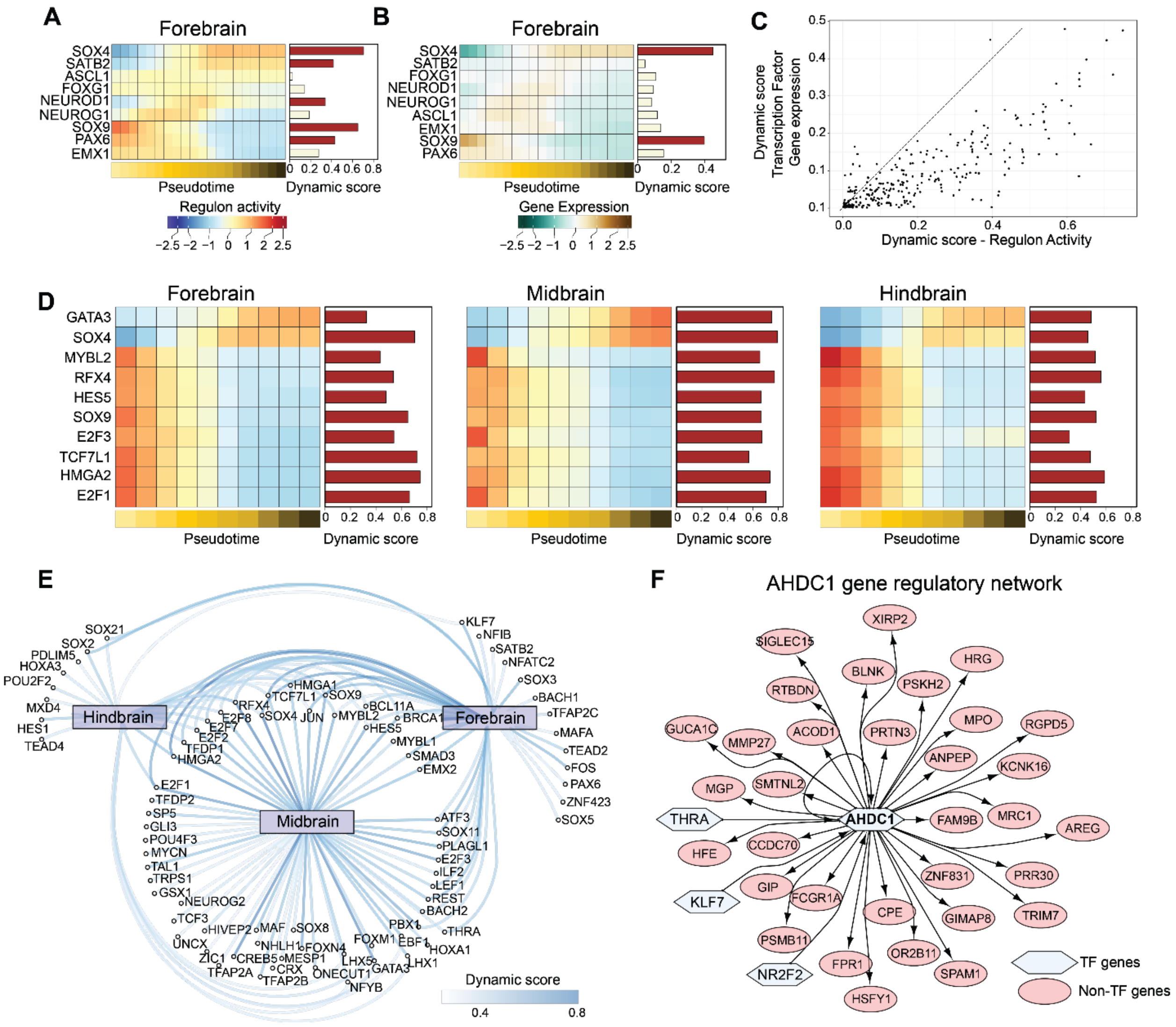
A
Forebrain
SOX4
SATB2
ASCL1
FOXG1
NEUROD1
NEUROG1
SOX9
PAX6
EMX1
0 0.2 0.4 0.6 0.8
Pseudotime
Dynamic score
Regulon activity
−2.5 −2 −1 0 1 2 2.5
B
Forebrain
SOX4
SATB2
FOXG1
NEUROD1
NEUROG1
ASCL1
EMX1
SOX9
PAX6
0 0.2 0.4
Pseudotime
Dynamic score
Gene Expression
−2.5 −2 −1 0 1 2 2.5
C
Dynamic score
Transcription Factor
Gene expression
0.5
0.4
0.3
0.2
0.1
0.1
0.0 0.2 0.4 0.6
Dynamic score - Regulon Activity
D
Forebrain
Midbrain
Hindbrain
GATA3
SOX4
MYBL2
RFX4
HES5
SOX9
E2F3
TCF7L1
HMGA2
E2F1
0 0.2 0.4 0.6 0.8
Pseudotime
Dynamic score
E
Hindbrain
Forebrain
Midbrain
SOX21
SOX2
PDLIM5
HOXA3
POU2F2
MXD4
HES1
TEAD4
KLF7
NFIB
SATB2
NFATC2
SOX3
BACH1
TFAP2C
MAFA
TEAD2
FOS
PAX6
ZNF423
SOX5
HMGA1
TCF7L1
SOX9
RFX4
E2F8
E2F7
E2F2
TFDP1
HMGA2
SOX4
JUN
MYBL2
BCL11A
BRCA1
HES5
MYBL1
SMAD3
EMX2
E2F1
TFDP2
SP5
GLI3
POU4F3
MYCN
TAL1
TRPS1
GSX1
NEUROG2
TCF3
UNCX
HIVEP2
MAF
SOX8
ZIC1
NHLH1
CREB5
MESP1
FOXN4
TFAP2A
CRX
TFAP2B
ONECUT1
NFYB
LHX5
GATA3
LHX1
FOXM1
EBF1
PBX1
HOXA1
THRA
ATF3
SOX11
PLAGL1
E2F3
ILF2
LEF1
REST
BACH2
Dynamic score
0.4
0.8
F
AHDC1 gene regulatory network
AHDC1
XIRP2
SIGLEC15
BLNK
PSKH2
HRG
RTBDN
MPO
RGPD5
GUCA1C
MMP27
ACOD1
PRTN3
ANPEP
KCNK16
MGP
SMTNL2
THRA
FAM9B
MRC1
AREG
HFE
CCDC70
ZNF831
PRR30
KLF7
GIP
FCGR1A
CPE
GIMAP8
TRIM7
PSMB11
OR2B11
FPR1
SPAM1
NR2F2
HSFY1
TF genes
Non-TF genes

**Figure S2: Integrative analysis of transcription factor dynamics and regulatory architecture across broad neurogenesis lineages.**

**A.** Binned pseudotemporal variation in regulon activity for transcription factors shown in **Fig. 2D**, for the human forebrain.

**B.** Pseudotemporal gene expression profiles for forebrain neurogenic transcription factors.

**C.** Scatter plot comparing dynamic scores derived from regulon activity and corresponding transcription factor gene expression. Statistical significance was assessed using a paired Wilcoxon signed-rank test (Wilcoxon signed-rank test, $p \leq 2.2e{-16}$).

**D.** Pseudotemporal regulon activity profiles for highly dynamic lineage-conserved regulons in the forebrain, midbrain, and hindbrain. Bar plots denote dynamic scores, with red indicating values greater than 0.3 and tan indicating less than 0.3.

**E.** Bipartite network visualization linking broad neurogenic lineages to their highly dynamic regulons. Connections with dynamic scores >0.4 are displayed.

**F.** Transcription factor–target gene regulatory network for AHDC1 in the human forebrain. Regulators are shown as hexagonal nodes, targets as elliptical nodes, with arrows indicating regulatory direction.

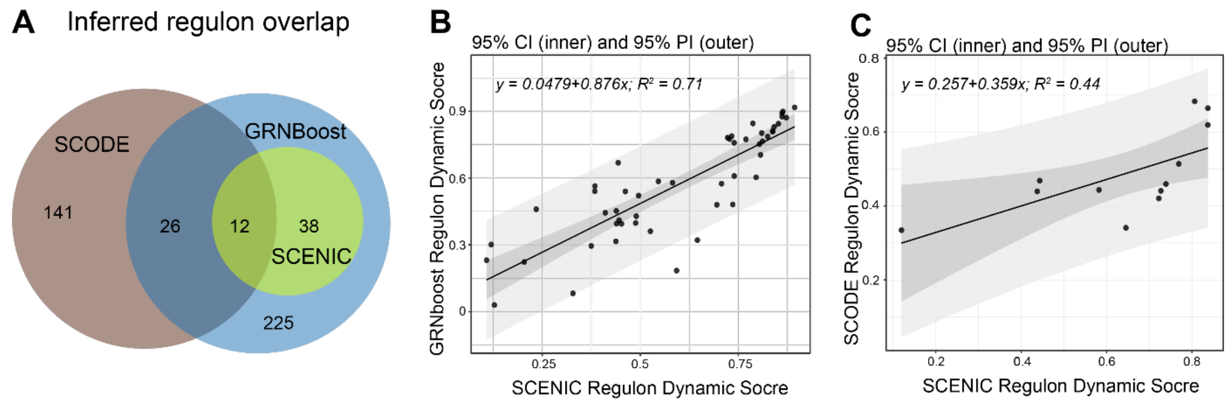


**Figure S3: Sensitivity analysis for gene regulatory network (GRN) inference methods.**

**A.** Venn diagram depicting the regulon overlap derived from three GRN inference methods: SCENIC[1,3], SCODE[4], GRNBoost[5]. The GRNs were inferred for the human telencephalon CGE neuronal lineage using the top 5000 highly variable genes. Transcription factors with more than 5 regulatory targets were defined as regulons.

**B-C**, scatter plotvisualizations for regulon dynamic scores for regulons derived from GRNBoost and SCENIC (**B**)and SCODE and SCENIC (**C**). Only the regulons overlapped in pair-wise manner were included in the scatter plots.

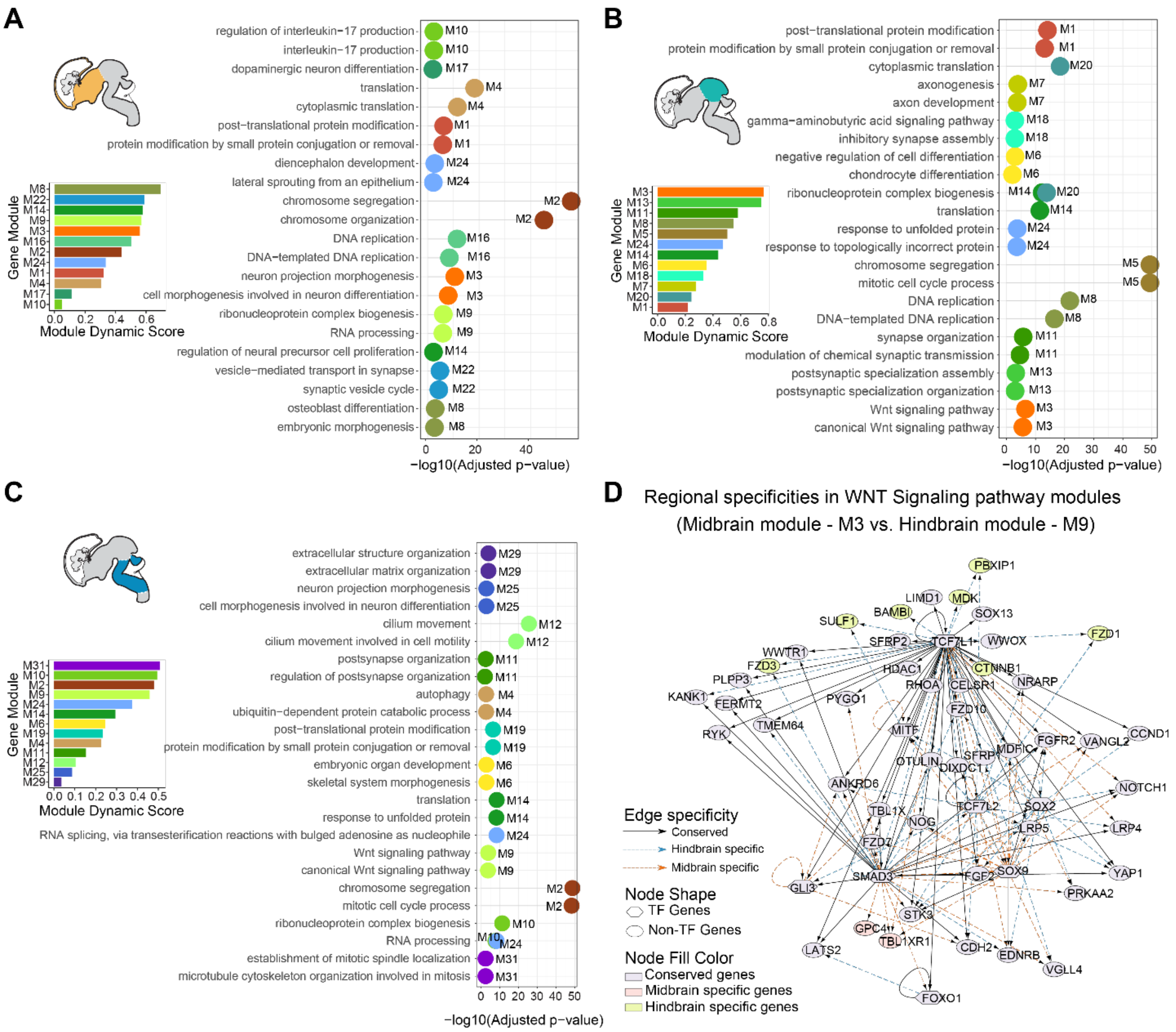
A
regulation of interleukin−17 production
interleukin−17 production
dopaminergic neuron differentiation
translation
cytoplasmic translation
post−translational protein modification
protein modification by small protein conjugation or removal
diencephalon development
lateral sprouting from an epithelium
chromosome segregation
chromosome organization
DNA replication
DNA−templated DNA replication
neuron projection morphogenesis
cell morphogenesis involved in neuron differentiation
ribonucleoprotein complex biogenesis
RNA processing
regulation of neural precursor cell proliferation
vesicle−mediated transport in synapse
synaptic vesicle cycle
osteoblast differentiation
embryonic morphogenesis
−log10(Adjusted p−value)
Gene Module
Module Dynamic Score
B
post−translational protein modification
protein modification by small protein conjugation or removal
cytoplasmic translation
axonogenesis
axon development
gamma−aminobutyric acid signaling pathway
inhibitory synapse assembly
negative regulation of cell differentiation
chondrocyte differentiation
ribonucleoprotein complex biogenesis
translation
response to unfolded protein
response to topologically incorrect protein
chromosome segregation
mitotic cell cycle process
DNA replication
DNA−templated DNA replication
synapse organization
modulation of chemical synaptic transmission
postsynaptic specialization assembly
postsynaptic specialization organization
Wnt signaling pathway
canonical Wnt signaling pathway
−log10(Adjusted p−value)
Gene Module
Module Dynamic Score
C
extracellular structure organization
extracellular matrix organization
neuron projection morphogenesis
cell morphogenesis involved in neuron differentiation
cilium movement
cilium movement involved in cell motility
postsynapse organization
regulation of postsynapse organization
autophagy
ubiquitin−dependent protein catabolic process
post−translational protein modification
protein modification by small protein conjugation or removal
embryonic organ development
skeletal system morphogenesis
translation
response to unfolded protein
RNA splicing, via transesterification reactions with bulged adenosine as nucleophile
Wnt signaling pathway
canonical Wnt signaling pathway
chromosome segregation
mitotic cell cycle process
ribonucleoprotein complex biogenesis
RNA processing
establishment of mitotic spindle localization
microtubule cytoskeleton organization involved in mitosis
−log10(Adjusted p−value)
Gene Module
Module Dynamic Score
D
Regional specificities in WNT Signaling pathway modules
(Midbrain module - M3 vs. Hindbrain module - M9)
Edge specificity
Conserved
Hindbrain specific
Midbrain specific
Node Shape
TF Genes
Non-TF Genes
Node Fill Color
Conserved genes
Midbrain specific genes
Hindbrain specific genes

**Figure S4: Co-regulatory gene module dynamics and their biological interpretations.**

**A–C.** Gene Ontology (GO): Biological Process terms enriched in co-regulatory gene modules for the forebrain (**A**), midbrain (**B**), and hindbrain (**C**). Inset bar plots display the corresponding dynamic module scores, derived from gene module activity estimates across corresponding neurogenesis lineages. Only modules with statistically significant ($p < 0.01$) GO: Biological Process enrichments are included.

**D.** Differential gene regulatory network visualization of region-specific regulatory relationships governing WNT signaling pathway–associated modules. Both midbrain module-M3 and hindbrain module-M9 are enriched for WNT signaling genes, identified by intersecting WNT pathway annotations with module memberships. Network nodes are colored by regional specificity. Transcription factor genes are represented as hexagons, non–transcription factor genes as ellipses. Regulatory interactions are visualized as arrows, with edge shapes denoting specificity within the forebrain and hindbrain GRNs.

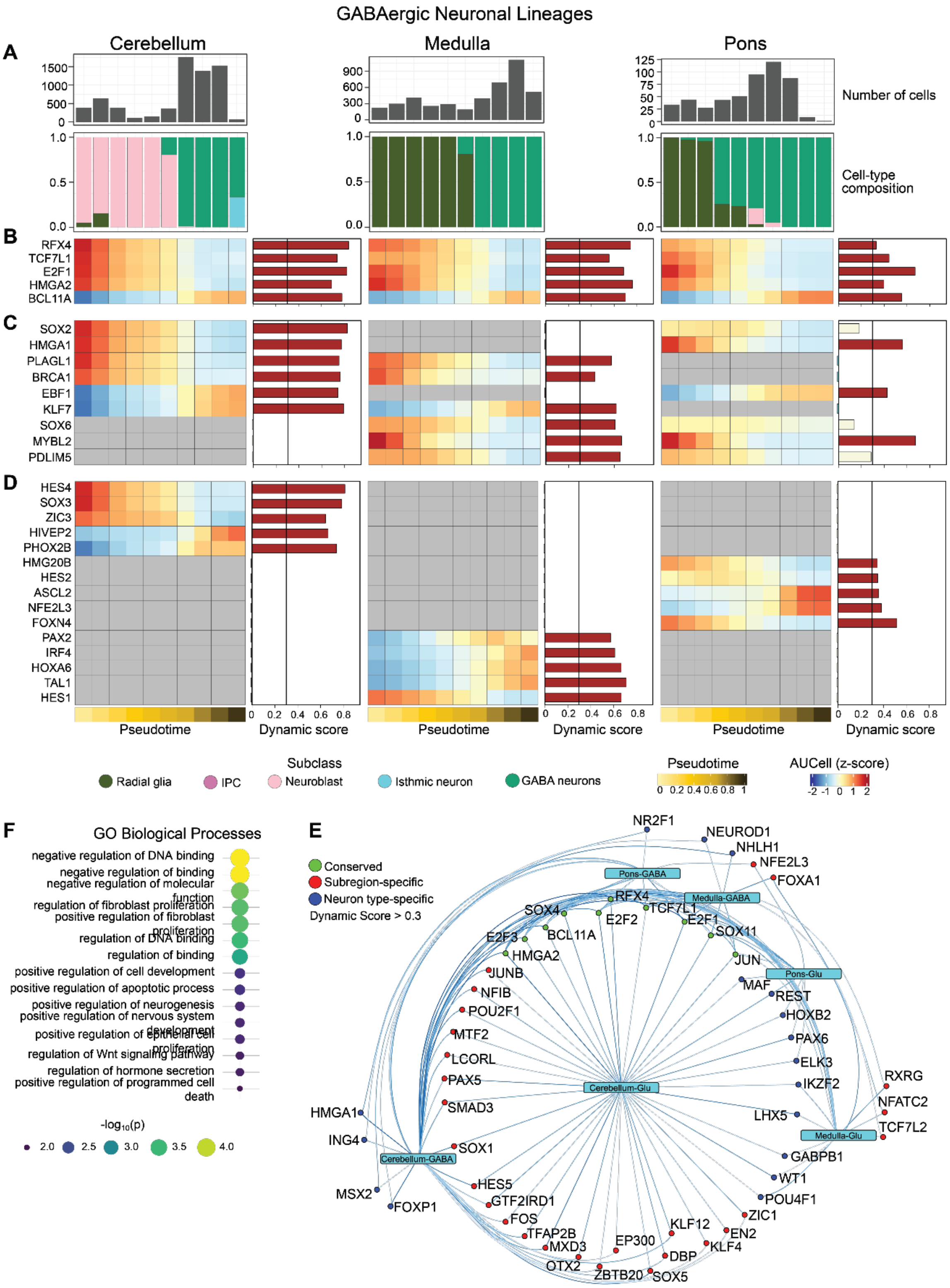

GABAergic Neuronal Lineages
A
Cerebellum
Medulla
Pons
Number of cells
Cell-type composition
B
RFX4
TCF7L1
E2F1
HMGA2
BCL11A
C
SOX2
HMGA1
PLAGL1
BRCA1
EBF1
KLF7
SOX6
MYBL2
PDLIM5
D
HES4
SOX3
ZIC3
HIVEP2
PHOX2B
HMG20B
HES2
ASCL2
NFE2L3
FOXN4
PAX2
IRF4
HOXA6
TAL1
HES1
Pseudotime
Dynamic score
Subclass
Radial glia
IPC
Neuroblast
Isthmic neuron
GABA neurons
Pseudotime
AUCell (z-score)
F
GO Biological Processes
negative regulation of DNA binding
negative regulation of binding
negative regulation of molecular function
regulation of fibroblast proliferation
positive regulation of fibroblast proliferation
regulation of DNA binding
regulation of binding
positive regulation of cell development
positive regulation of apoptotic process
positive regulation of neurogenesis
positive regulation of nervous system development
positive regulation of epithelial cell proliferation
regulation of Wnt signaling pathway
regulation of hormone secretion
positive regulation of programmed cell death
-log10(p)
E
Conserved
Subregion-specific
Neuron type-specific
Dynamic Score > 0.3
Pons-GABA
Medulla-GABA
Pons-Glu
Cerebellum-Glu
Medulla-Glu
Cerebellum-GABA
NR2F1
NEUROD1
NHLH1
NFE2L3
FOXA1
RFX4
SOX4
TCF7L1
E2F2
E2F1
E2F3
BCL11A
SOX11
HMGA2
JUN
JUNB
MAF
NFIB
REST
POU2F1
HOXB2
MTF2
PAX6
LCORL
ELK3
PAX5
IKZF2
RXRG
SMAD3
NFATC2
HMGA1
LHX5
TCF7L2
ING4
SOX1
GABPB1
MSX2
FOXP1
HES5
WT1
GTF2IRD1
POU4F1
FOS
ZIC1
TFAP2B
KLF12
EN2
MXD3
EP300
KLF4
OTX2
DBP
ZBTB20
SOX5


96

**Figure S5. Regulon dynamics across human hindbrain subregions—GABAergic neuronal lineages.**

**A.** Bar plots showing the number of cells (top) and cell type composition colored by cell type (bottom) across pseudotime bins for cerebellar, medullary, and pontine GABAergic neuronal lineages. RG, radial glia; IPC, intermediate progenitor cells

**B–D.** Pseudotemporal profiles of regulon activity z-scores of selected regulons classified as: conserved across all three lineages and highly dynamic (**B**), pairwise conserved (i.e., present in two of three lineages; **C**), and uniquely active in one lineage (**D**). Bar colors indicate dynamic score, with red representing scores > 0.3 and tan < 0.3.

**E**. Bipartite network linking each lineage to dynamic regulons featured in the UpSet plot in **Figure 3E**. Green denotes regulons conserved across all six lineages; red highlights those specific to a hindbrain subregion regardless of neuronal identity; and blue indicates regulons associated with a neuronal type.

**F**. Dot plot showing enriched Gene Ontology (GO: Biological Processes) terms for regulons conserved across all six lineages.

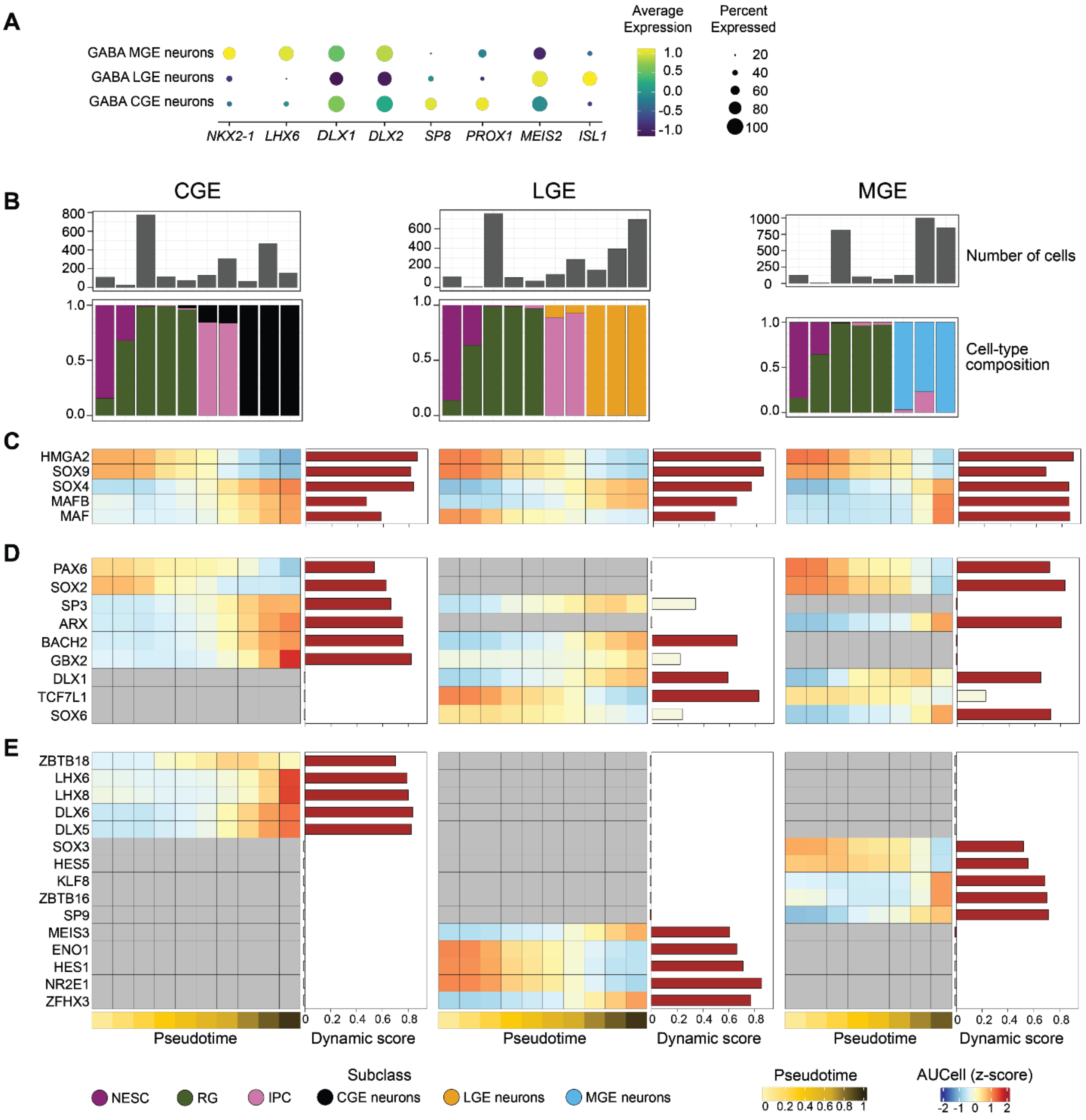
A
Average Expression
Percent Expressed
GABA MGE neurons
GABA LGE neurons
GABA CGE neurons
NKX2-1
LHX6
DLX1
DLX2
SP8
PROX1
MEIS2
ISL1
1.0
0.5
0.0
-0.5
-1.0
20
40
60
80
100
B
CGE
LGE
MGE
Number of cells
Cell-type composition
C
HMGA2
SOX9
SOX4
MAFB
MAF
D
PAX6
SOX2
SP3
ARX
BACH2
GBX2
DLX1
TCF7L1
SOX6
E
ZBTB18
LHX6
LHX8
DLX6
DLX5
SOX3
HES5
KLF8
ZBTB16
SP9
MEIS3
ENO1
HES1
NR2E1
ZFHX3
Pseudotime
Dynamic score
Subclass
NESC
RG
IPC
CGE neurons
LGE neurons
MGE neurons
Pseudotime
0 0.2 0.4 0.6 0.8 1
AUCell (z-score)
-2 -1 0 1 2

**Figure S6. Regulon dynamics across telencephalic GABAergic neuronal subtypes.**

**A.** Dot plot showing average expression of subtype-specific markers across CGE-, LGE-, and MGE-derived neurons.

**B.** Bar plots depicting the number of cells (top) and lineage-specific cell type composition (bottom) across pseudotime bins for CGE, LGE, and MGE lineages. CGE, caudal ganglionic eminence; MGE, medial ganglioic eminence; LGE, lateral ganglionic eminence

**C–E.** Pseudotemporal profiles of regulon activity z-scores of representative regulons grouped by conservation patterns: conserved and dynamically active in all three lineages (**C**), pairwise conserved (present in two of three lineages) (**D**) and unique to a single lineage (**E**). Bar colors indicate dynamic score: red (> 0.3) and tan (< 0.3).

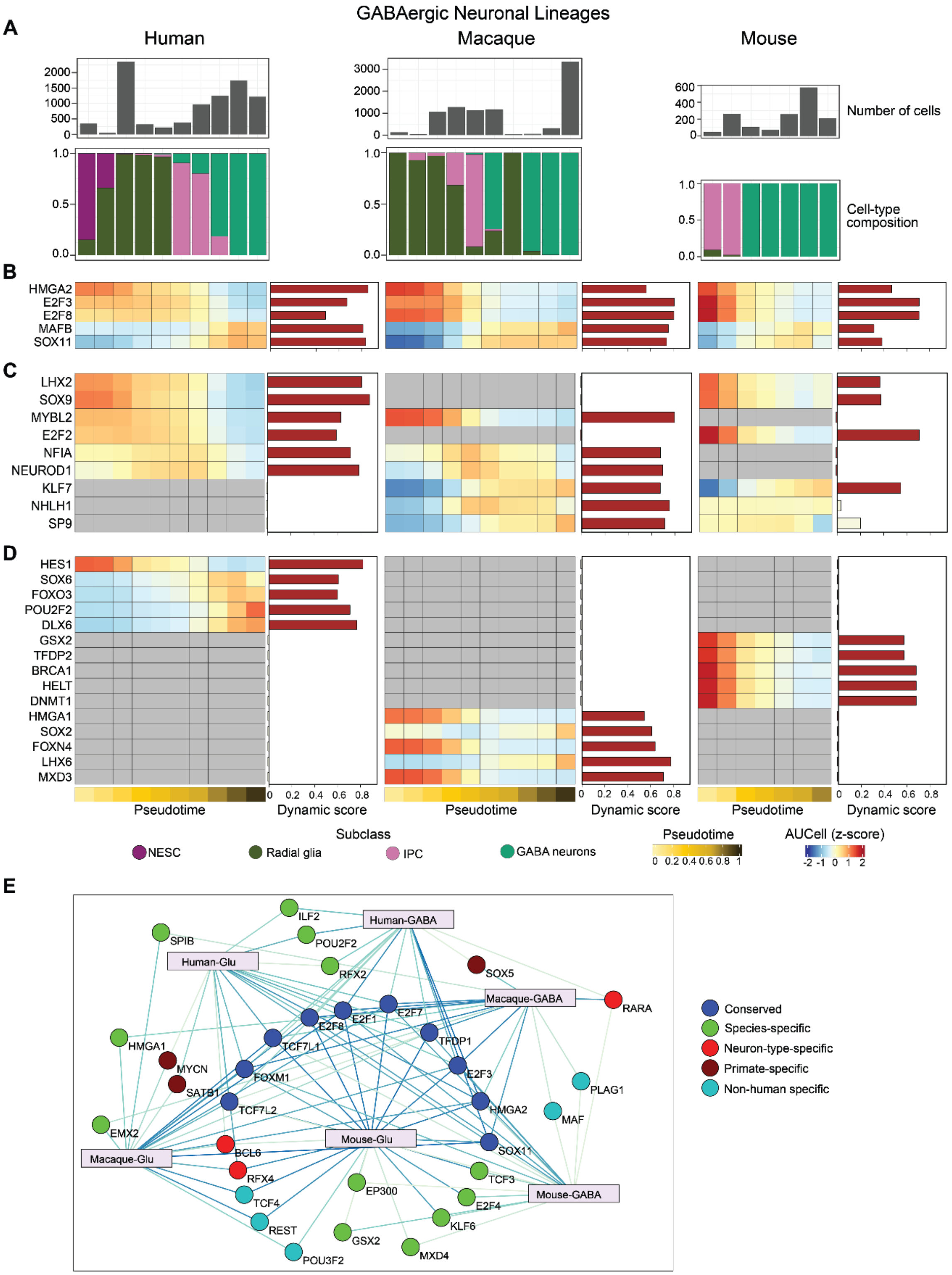
GABAergic Neuronal Lineages
A
Human
Macaque
Mouse
Number of cells
Cell-type composition
B
HMGA2
E2F3
E2F8
MAFB
SOX11
C
LHX2
SOX9
MYBL2
E2F2
NFIA
NEUROD1
KLF7
NHLH1
SP9
D
HES1
SOX6
FOXO3
POU2F2
DLX6
GSX2
TFDP2
BRCA1
HELT
DNMT1
HMGA1
SOX2
FOXN4
LHX6
MXD3
Pseudotime
Dynamic score
Subclass
NESC
Radial glia
IPC
GABA neurons
AUCell (z-score)
E
Human-GABA
Human-Glu
Macaque-GABA
Macaque-Glu
Mouse-Glu
Mouse-GABA
Conserved
Species-specific
Neuron-type-specific
Primate-specific
Non-human specific

**Figure S7: Cross-species regulon dynamics in telencephalic GABAergic neurogenesis.**

**A.** Bar plots showing the number of cells (top) and cell type composition (bottom) across pseudotime bins for human, macaque, and mouse GABAergic lineages. NESC, neuroepithelial stem cell; RG, radial glia; IPC, intermediate progenitor cell.

**B–D.** Pseudotemporal activity profiles of selected regulons grouped by conservation: conserved and highly dynamic across all three species (**B**), pairwise conserved (present in two of three species, **C**), and unique to individual species (**D**). Bar colors denote dynamic scores: red (> 0.3) and tan (< 0.3).

**E.** Bipartite network linking each lineage to dynamic regulons featured in the UpSet plot in **Figure 6E**. Color codes: blue - conserved across all six glutamatergic lineages, green - species-specific, red - neuron-type-specific, brown - primate-specific and cyan - non-human-specific.

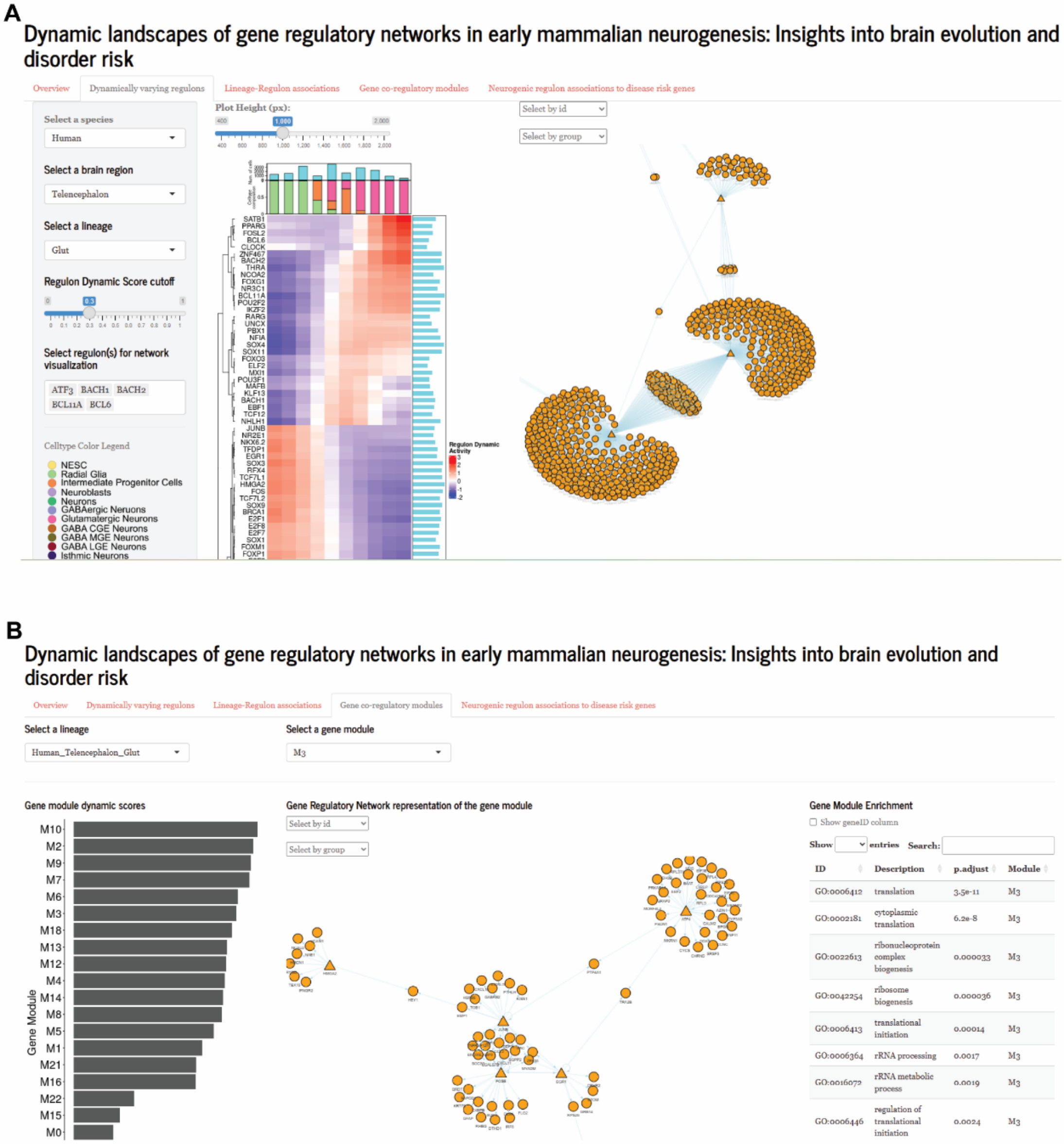
A
Dynamic landscapes of gene regulatory networks in early mammalian neurogenesis: Insights into brain evolution and disorder risk
Overview
Dynamically varying regulons
Lineage-Regulon associations
Gene co-regulatory modules
Neurogenic regulon associations to disease risk genes
Select a species
Human
Select a brain region
Telencephalon
Select a lineage
Glut
Regulon Dynamic Score cutoff
Select regulon(s) for network visualization
ATF3 BACH1 BACH2 BCL11A BCL6
Celltype Color Legend
NESC
Radial Glia
Intermediate Progenitor Cells
Neuroblasts
Neurons
GABAergic Neurons
Glutamatergic Neurons
GABA CGE Neurons
GABA MGE Neurons
GABA LGE Neurons
Isthmic Neurons
Plot Height (px):
Select by id
Select by group
Regulon Dynamic Activity
B
Dynamic landscapes of gene regulatory networks in early mammalian neurogenesis: Insights into brain evolution and disorder risk
Overview
Dynamically varying regulons
Lineage-Regulon associations
Gene co-regulatory modules
Neurogenic regulon associations to disease risk genes
Select a lineage
Human_Telencephalon_Glut
Select a gene module
M3
Gene module dynamic scores
Gene Module
M10
M2
M9
M7
M6
M3
M18
M13
M12
M4
M14
M8
M5
M1
M21
M16
M22
M15
M0
Gene Regulatory Network representation of the gene module
Select by id
Select by group
Gene Module Enrichment
Show geneID column
Show entries Search:
ID Description p.adjust Module
GO:0006412 translation 3.5e-11 M3
GO:0002181 cytoplasmic translation 6.2e-8 M3
GO:0022613 ribonucleoprotein complex biogenesis 0.000033 M3
GO:0042254 ribosome biogenesis 0.000036 M3
GO:0006413 translational initiation 0.00014 M3
GO:0006364 rRNA processing 0.0017 M3
GO:0016072 rRNA metabolic process 0.0019 M3
GO:0006446 regulation of translational initiation 0.0024 M3

**Figure S8: Interactive exploration of dynamic neurogenic regulatory networks.**

**A.** Exploration of dynamic regulons for a selected for neurogenic lineage. A user can specify the species, brain region and a lineage and the webapp will provide a heatmap of all the dynamic regulons depicting their temporal activity. User can also select any number of regulons from a drop-down menu to explore their regulatory interactions.

**B.** Exploration of dynamic co-regulatory gene modules for a selected human neurogenic lineage. The user can specify the lineage and it will provide a bar plot ranking the gene module based on their dynamic behavior. The user can also select a co-regulatory gene module for network visualization and to explore their functional enrichment.
